# Equivalent volitional learning emerges through circuit-specific population dynamics in motor cortex and hippocampus

**DOI:** 10.64898/2026.06.04.730137

**Authors:** Andres de Vicente, Catalin Mitelut, Renan V. Mendes, Lorenzo Marianelli, Mariona Colomer Rosell, David Bruckner, Giampiero Bardella, Flavio Donato

## Abstract

Learning operates across different brain circuits to associate population activity patterns with desired outcomes, and to enable volitional reactivation of those patterns to control behavior. These circuits differ profoundly in their architecture and dynamical regimes, yet which features of learning are shared across them and which arise from circuit-specific implementations remains unknown. Here, we use a brain-computer interface (BCI) to train mice to modulate the activity of selected neuronal ensembles toward configurations that trigger reward delivery. By making reward delivery contingent directly on population activity, we impose an identical associative learning problem on two circuits with distinct dynamical regimes: the primary motor cortex (M1) and the hippocampal area CA3. Mice acquired robust volitional control in both regions, and learning produced a set of shared signatures across circuits, including modulation of reward-controlling neurons, network-level sparsification, and greater exploration of reward-related activity patterns. These signatures were underpinned by distinct population dynamics: M1 activity flowed continuously through reward-associated states, whereas CA3 activity traced approach-and-return dynamics around them. Recurrent network models endowed with distinct minimal connectivity constraints chosen to reflect the dominant dynamical regime associated with each region captured key features of these shared signatures and region-specific dynamics, indicating that local architectural constraints are sufficient to account for the distinct implementations of learning. These findings indicate that equivalent learning outcomes arise from divergent dynamical implementations across architecturally distinct circuits. This principled degeneracy reveals that learning is not a single canonical solution, but is implemented through multiple circuit-specific mechanisms shaped by local network architecture.

## Introduction

Associative learning is at once widespread and specialized across the brain: many circuits can encode new associations, yet each underpins a distinct set of behavioral skills, from executing complex motor sequences to navigating spatial environments^1–4^. Across circuits, learning operates through two linked processes: associating population activity patterns with desired outcomes, and acquiring the ability to volitionally reactivate those patterns to drive behavior^5–9^. The circuits that support these operations differ profoundly in their architecture, connectivity, and dynamical regimes^10–12^, yet which features of learning are shared across them and which arise from circuit-specific implementations remains unknown.

Degeneracy, the principle that equivalent functional outcomes can arise through multiple mechanistic implementations^13–18^, predicts that different circuits satisfy the same learning demands through distinct dynamical mechanisms shaped by circuit architecture. Testing this prediction experimentally has been difficult because circuits that differ most in architecture also typically support distinct behavioral functions, making it impossible to determine whether differences in learning-related dynamics reflect intrinsic circuit properties or behavior-specific demands. Brain-computer interface (BCI) paradigms offer a principled solution^19–25^. By making reward contingent directly on neuronal population activity, the same associative learning problem can be imposed across circuits while holding behavioral demands constant, thereby isolating the influence of circuit architecture and its associated dynamical constraints.

The primary motor cortex (M1) and hippocampal area CA3 provide an ideal contrast for testing this prediction. Both regions share key structural features, including dense recurrent connectivity and robust synaptic plasticity^26–30^. Despite these similarities, they are commonly described through distinct dominant computational regimes. M1 population dynamics are organized as structured trajectories through a low-dimensional neural state space that support the generation of precisely timed motor outputs^12,31–36^. CA3 is classically conceptualized as an attractor-based autoassociative network supporting memory storage and retrieval through pattern completion^26,37–41^. If local dynamical constraints shape how learning is implemented, these two regions should achieve equivalent learning outcomes through distinct dynamical mechanisms.

Using longitudinal two-photon calcium imaging during BCI learning, we compared how M1 and CA3 populations solve the same volitional learning problem: controlling the activity of specific neuronal ensembles to obtain reward^20,24,42–46^. Mice acquired robust volitional control when BCI learning was directed to either region, and exhibited shared network-level signatures of learning, including selective modulation of reward-controlling neurons, sparsification of reward-related activity, and increased exploration of reward-associated population states. Despite these shared outcomes, the underlying population dynamics diverged distinctively between regions: within trials, M1 activity progressed as continuous trajectories through reward-associated states, whereas CA3 activity developed stereotyped approach-and-return dynamics centered around them. Recurrent network models endowed with distinct minimal constraints and trained in an *in silico* version of the BCI task captured key features of these shared and divergent solutions while converging on equivalent task performance.

Together, these results indicate that equivalent volitional learning is achieved through circuit-specific dynamical implementations. This principled degeneracy, where shared learning signatures emerge alongside distinct population dynamics in both biological circuits and recurrent network models, reveals that architectural constraints shape how learning is implemented across circuits even under equivalent task demands.

### Equivalent volitional learning emerges in M1 and CA3 under identical task demands

To determine whether distinct recurrent circuits solve the same learning problem through shared or circuit-specific dynamics, we trained mice on an identical closed-loop BCI task (adapted from Clancy et al., 2014^20^) targeted either to primary motor cortex (M1) or hippocampal area CA3. In this task, mice learned to volitionally modulate the combined activity of two distinct neuronal ensembles, each comprising two neurons, to obtain rewards. The activity of the two ensembles were combined online into a single ensemble state, defined as the sum of the activity of neurons in one ensemble (the “positive” ensemble) minus the sum of the activity of neurons in the other (the “negative” ensemble). Ensemble state was translated in real time into an auditory feedback tone (the sound cursor) whose frequency tracked the ensemble state value: activity in the positive ensemble raised the cursor frequency, while activity in the negative ensemble lowered it (Figure 1A, B). A liquid sucrose reward was delivered when the cursor crossed a calibrated frequency threshold, corresponding to a target ensemble state value. Two-photon calcium imaging in head-fixed mice monitored activity in both the positive and negative ensemble neurons (the direct neurons), as well as in the rest of the network (the indirect neurons). Real-time computation of ensemble state transformed neuronal activity into the auditory feedback signal via Open-CaBCI (Extended Data Figure 1A), an open-source Python framework we developed for real-time calcium-imaging-based brain-computer interface tasks (see Methods). The BCI was independently targeted to either the primary motor cortex (M1) or the hippocampal area CA3 across two separate cohorts of mice, with ensemble activity monitored through a cranial window or a microendoscope, respectively (Figure 1A and Methods. M1 cohort: 30388 neurons recorded across 7 animals over 54 sessions; 3995 unique neurons were longitudinally tracked through training. CA3 cohort: 23168 neurons across 6 animals over 48 sessions; 2773 unique neurons were longitudinally tracked through training^47,48^).

**Figure 1.**
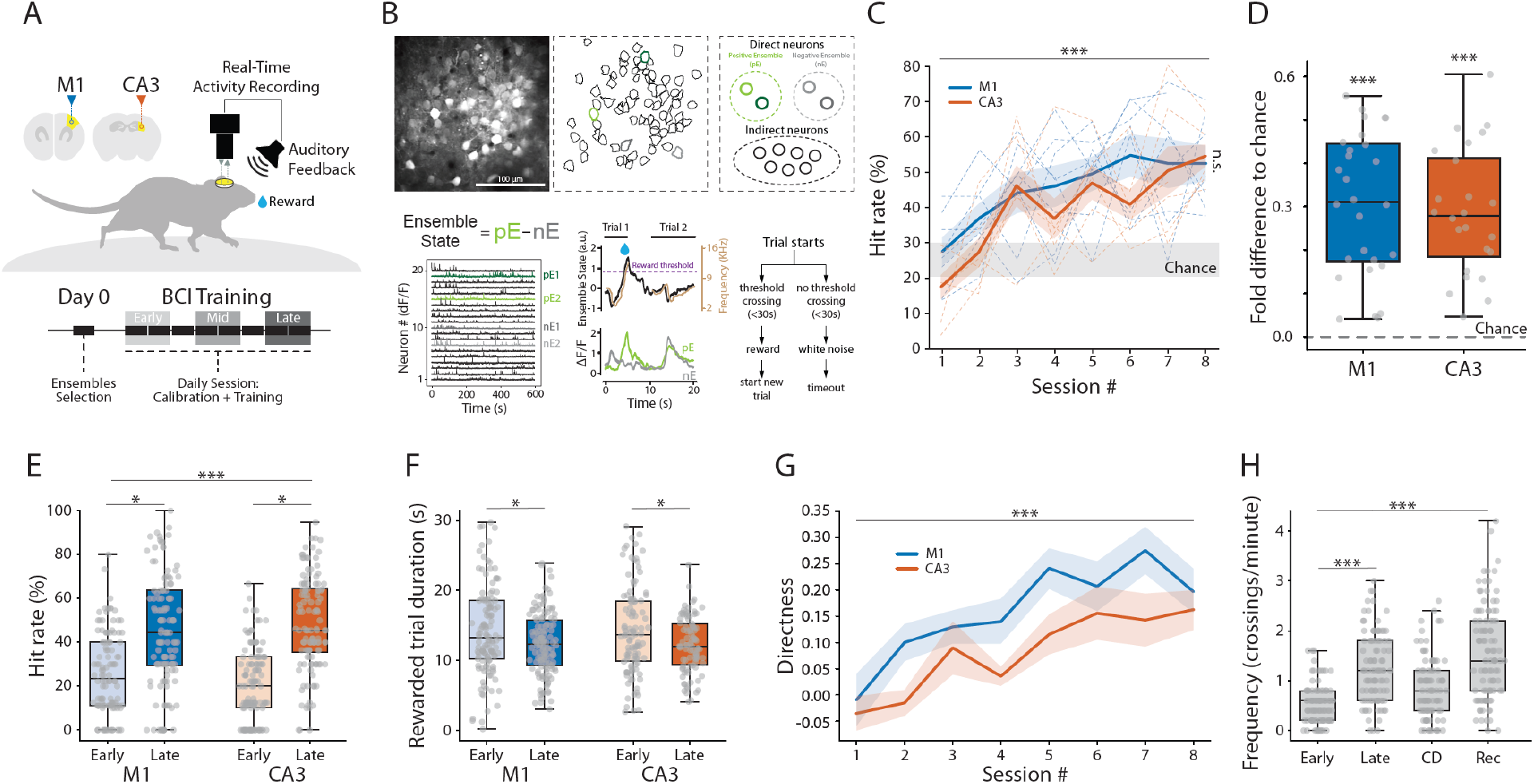
Equivalent volitional BCI learning in M1 and CA3. **A**, Schematic of the closed-loop two-photon calcium imaging BCI paradigm. Head-fixed mice were trained to volitionally modulate selected neuronal ensembles in M1 or CA3. The protocol comprised an ensemble-selection session (day 0) followed by 8 daily training sessions, each with a 15-min calibration period and a 50-min BCI session. Throughout the manuscript, Early, Mid, and Late learning refer to sessions 1–2, 4–5, and 7–8, respectively. **B**, Example field of view and task structure. Top left, representative imaging field with labeled neurons and the BCI ensemble subset. Top right, ensemble composition: two neurons assigned to the direct positive ensemble (pE; shades of green), two to the direct negative ensemble (nE; shades of gray), with remaining recorded neurons as indirect. Bottom left, example calcium traces from the same session (green, positive; gray, negative). Bottom middle, ensemble state computed online as (pE1 + pE2) − (nE1 + nE2). Ensemble state is converted into a sound cursor whose frequency tracks the state value (brown). Reward was delivered when the cursor crosses a calibrated threshold. Bottom right, trial structure: up to 30-s response window; threshold crossing triggered reward and a new trial; failure triggered white-noise feedback followed by a timeout. **C**, Longitudinal task performance across training in M1 (blue, *n* = 7 mice) and CA3 (orange, *n* = 6 mice) cohorts. Thin dashed lines, individual animals; thick lines, cohort mean ± s.e.m. Gray shading, chance-level performance. Both cohorts showed progressive improvement across sessions. Linear mixed-effects model: significant effect of session (β = 0.045, *p* < 0.001); no main effect of group (β = 0.054, *p* = 0.118) or group × session interaction (β = −0.008, *p* = 0.573). Session-wise comparisons not significant after FDR correction. Animal-level integrated performance did not differ between groups (two-sided Mann–Whitney, *p* = 0.138). **D**, Session-level above-chance performance during late learning (task hit rate minus matched chance hit rate) for M1 and CA3. Gray points, individual sessions; boxplots, median and interquartile range; whiskers, data range; dashed line, chance. Both regions showed positive late-phase differences (M1: median Δ = 0.311, *n* = 28 sessions, one-sided Wilcoxon signed-rank vs zero, *p* < 0.001; CA3: median Δ = 0.279, *n* = 24 sessions, *p* < 0.001). No significant difference between regions (two-sided Mann–Whitney, *p* = 0.666). **E**, Early versus late training task performance in M1 and CA3. Boxplots, pooled 5-min hit-rate bins; overlaid points, individual bins. Mixed-effects model accounting for repeated measurements across mice and sessions: significant increase from early to late (β = 0.867, *p* < 0.001); no group × phase interaction (β = −0.208, *p* = 0.279). As secondary post hoc analyses, group-specific early-versus-late comparisons remained significant after Benjamini–Hochberg false-discovery-rate correction in both M1 (FDR-adjusted *p* = 0.031) and CA3 (FDR-adjusted *p* = 0.031), indicating improved performance in both groups. **F**, Rewarded-trial duration across early and late phases of learning in M1 and CA3. Boxplots, chunk-level (5-min) rewarded-trial durations; overlaid points, individual chunks. Mixed-effects model accounting for repeated measurements across mice and sessions: significant reduction from early to late (β = −2.423, *p* = 0.017); no group × phase interaction (β = 0.699, *p* = 0.606). Group-specific contrasts (BH-corrected): M1 β = −1.752, FDR-adjusted *p* = 0.031; CA3 β = −2.390, FDR-adjusted *p* = 0.041. **G**, Directness of auditory-feedback trajectories across training in M1 (blue) and CA3 (orange). Directness, net trajectory change divided by cumulative absolute stepwise change: values near 1, monotonic and goal-directed; near 0, back-and-forth fluctuation; negative, net movement opposite the target. Lines, mean ± s.e.m. across sessions. Linear mixed-effects model: significant increase over training (β = 0.029, *p* < 0.001); higher directness in M1 than CA3 (β = 0.082, *p* = 0.002); no group × session interaction (β = 0.010, *p* = 0.339). **H**, Threshold-crossing rate across learning (Early, Late), contingency degradation (CD), and recovery (Rec) phases, in 5-min bins. Linear mixed-effects model. Relative to Early: Late β = 0.465, *p* < 0.001; Rec β = 0.942, *p* < 0.001; CD β = 0.249, *p* = 0.074. Benjamini–Hochberg-corrected pairwise contrasts: Early vs Late *p* < 0.001; Early vs Rec *p* < 0.001; Late vs Rec *p* < 0.001; CD vs Rec *p* < 0.001; Early vs CD and Late vs CD not significant. Selected comparisons are shown.

BCI training comprised an ensemble selection day (Day 0) followed by eight daily sessions (Figure 1A, B). On Day 0, neurons were selected and assigned to positive or negative ensembles (see Methods for the selection criteria); ensemble identity and imaging parameters were held fixed across all sessions to enable stable longitudinal tracking. At the start of each daily training session, the reward threshold was recalibrated based on spontaneous ensemble activity measured during a baseline recording period, targeting an expected success rate of 20–30% to maintain consistent task difficulty throughout training while accounting for the ensemble’s representational drift across days. Each session consisted of trial-based 30-second response windows, scored as successful if the ensemble state crossed the reward threshold (Figure 1B). A post-reward reset criterion required the ensemble state to return to below 50% of the threshold before the next trial started, preventing re-triggering due to residual elevated activity and ensuring post-reward dynamics were standardized across sessions and animals. Unsuccessful trials triggered a timeout before the next trial began. To enable direct comparison of learning trajectories across cohorts, training sessions were segmented into three epochs: early (sessions 1–2), mid (sessions 4–5), and late (sessions 7–8) learning.

In both M1 and CA3 cohorts, mice successfully learned to volitionally control neuronal activity to perform the BCI task. Rewarded trial percentage increased significantly across sessions from near-chance levels on day 1, with equivalent acquisition dynamics across regions (Figure 1C-E). Improvements accrued primarily across sessions rather than within individual training days (Extended Data Figure 1B). Trial duration decreased with learning (Figure 1F), and the sound cursor trajectory became more directed toward the reward threshold (Figure 1G), reflecting progressive refinement of volitional control across sessions. To assess whether task performance was contingent on a specific behavioral state, trials were classified as movement or stationary based on locomotion preceding reward delivery (Extended Data Figure 1C). Rewarded trials occurred in comparable proportions across movement and stationary trials (Extended Data Figure 1D), indicating that neither was preferentially associated with task success and that volitional control generalized across behavioral states.

To determine whether BCI performance reflected goal-directed volitional control of neural activity, or whether it reflected habitual neuronal activation through repeated reinforcement^7,24,49,50^, we performed contingency degradation experiments in a subset of M1 and CA3 animals. During these trials, reward delivery was decoupled from ensemble state by randomizing its timing, while threshold crossings triggered only the trial reset (as in Clancy et al., 2014)^26^. Disrupting the learned action-outcome association significantly reduced threshold crossing frequency and cursor directedness (Figure 1H; Extended Data Figure 1E), indicating that both success rate and performance efficiency were contingency-dependent. Recovery sessions reinstating the original closed-loop mapping restored both measures to pre-degradation levels (Figure 1H; Extended Data Figure 1E), confirming that volitional control of ensemble activity in both regions was goal-directed rather than habitual.

Together, these results indicate that M1 and CA3 both support robust volitional BCI learning under identical task demands, enabling a direct comparison of the population dynamics underlying learning in each circuit.

### Circuit-specific ensemble control strategies emerge during BCI learning

BCI learning requires animals to discover two linked features of the action-outcome contingency: which neurons drive reward delivery, and what activity patterns do so^5,6,9,23,51^. To track how these processes unfold over learning, we monitored mean firing rates of the positive and negative ensemble neurons across sessions (Figure 2A, B). In both M1 and CA3, ensemble neurons showed significantly elevated activity during task performance relative to calibration from the earliest training sessions, in both movement and stationary trials (Extended Data Figure 2A–C), which were therefore pooled for subsequent analyses. Calibration-period activity rates remained stable across days, indicating that these changes were specific to task performance rather than global drift. Ensemble engagement was thus established rapidly and maintained throughout learning, with ensemble state trajectories at threshold crossing remaining consistent across sessions in both regions (Extended Data Figure 2D). The relative engagement of positive and negative ensemble neurons evolved progressively. Early in learning, negative ensemble neurons exhibited significantly higher activity rates than positive ensemble neurons in both regions (Figure 2C). With learning, this asymmetry resolved differently across areas: in M1 it diminished toward symmetric engagement of both ensembles, whereas in CA3 it reversed, with positive ensemble neurons becoming significantly more active than negative ones by late learning (Figure 2C). No systematic within-session increase in ensemble activity was observed (Extended Data Figure 2E), consistent with performance improvements accumulating across rather than within sessions. The directness of ensemble state trajectories toward the reward threshold increased progressively with learning in both regions (Figure 2D), indicating that animals learned not only which activity patterns drive reward but also how to reach them more efficiently. During contingency degradation, task-associated ensemble activity modulation was reduced toward calibration baseline levels (Figure 2E), mirroring the behavioral sensitivity to action-outcome contingency.

**Figure 2.**
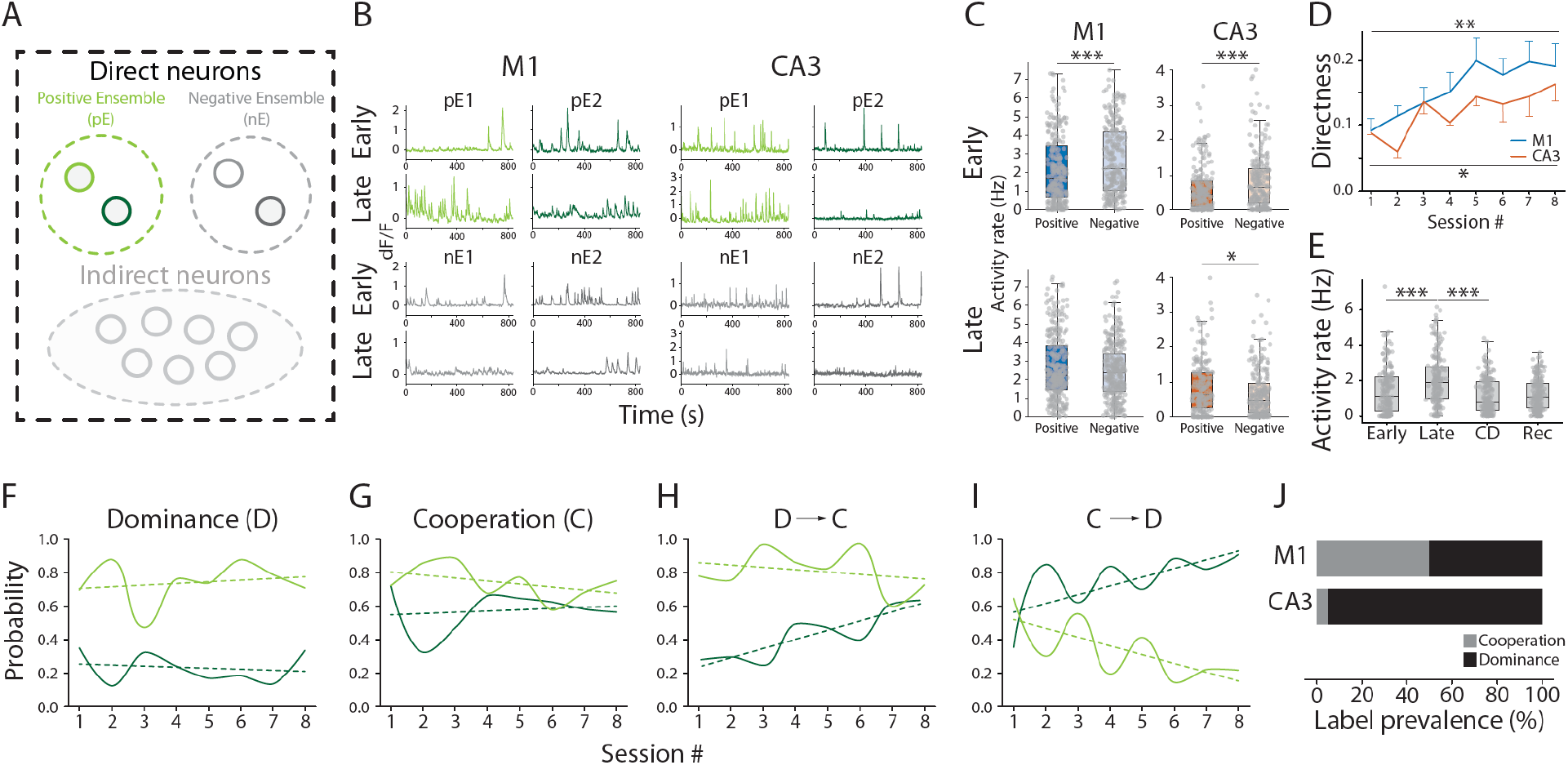
M1 and CA3 adopt distinct ensemble-control strategies during BCI learning. **A**, Schematic of BCI ensemble architecture, with direct neurons (i.e. positive and negative ensembles) highlighted. **B**, Example activity traces of BCI ensemble neurons in M1 and CA3 during early and late learning. Representative calcium traces are shown for the direct positive (pE1, pE2; green tones) and direct negative (nE1, nE2; gray tones) ensemble neurons. Upper rows, early learning; lower rows, late learning. **C**, Activity rates of positive and negative ensemble neurons during calibration and BCI-session epochs in M1 and CA3. Boxplots, 5-min bin values with overlaid datapoints. Linear mixed-effects models with class label (positive versus negative) as a fixed effect, accounting for repeated measurements across animals, with Benjamini–Hochberg correction across panel-wise tests. In the session period, negative ensemble neurons showed higher firing rates than positive ensemble neurons in early learning in both M1 (β_{NEG−POS} = 0.511, FDR-adjusted *p* < 0.001) and CA3 (β_{NEG−POS} = 0.205, FDR-adjusted *p* = 0.003). In late learning, this difference was not significant in M1 (β_{NEG−POS} = −0.130, FDR-adjusted *p* = 0.196); in CA3 the sign reversed, with positive ensemble neurons exhibiting higher firing rates than negative ensemble neurons (β_{NEG−POS} = −0.158, FDR-adjusted *p* = 0.014). **D**, Ensemble-trajectory directness across training sessions in M1 (blue) and CA3 (orange). Directness was computed on ensemble state and averaged per session. Lines, group mean ± s.d. across sessions. Linear regression on session-wise group means: both groups showed significant increases in directness across training (M1 slope = 0.016, *p* = 0.001; CA3 slope = 0.012, *p* = 0.012). Between-group comparison by permutation test on session-wise trajectories, with the L2 norm of the M1-versus-CA3 difference curve as test statistic: *p* = 0.341. **E**, Activity rate of positive ensemble neurons across learning, contingency degradation, and recovery phases. Boxplots, 5-min bin values across mice; overlaid points, individual bins. Linear mixed-effects model with stage as a fixed effect, accounting for repeated measurements across mice, with Benjamini–Hochberg correction across the six pairwise stage contrasts. All stage comparisons significant except CD versus Rec (early vs late, FDR-adjusted *p* < 0.001; early vs CD, FDR-adjusted *p* < 0.001; early vs Rec, FDR-adjusted *p* = 0.018; late vs CD, FDR-adjusted *p* < 0.001; late vs Rec, FDR-adjusted *p* < 0.001; CD vs Rec, FDR-adjusted *p* = 0.245). Selected comparisons are shown. **F–I**, Example reward-tuning trajectories of ensemble-positive neurons across training sessions. Solid lines, session-by-session evolution of reward-aligned burst probability for the two positive ensemble neurons in representative animals; dashed lines, corresponding linear trend fits. Each panel illustrates a distinct motif: dominance (D), one positive neuron shows persistently stronger reward tuning than the other; cooperation (C), both positive neurons show comparable reward tuning; D→C, initially dominant tuning becomes more cooperative across sessions; C→D, initially cooperative tuning becomes increasingly asymmetric. **J**, Prevalence of simplified reward-tuning labels in M1 and CA3. Configurations grouped into cooperation (both positive ensemble neurons contribute to reward-related activity) and dominance (reward-related activity primarily driven by one positive ensemble neuron). Stacked bars, percentage of configurations per category. Cooperation more prevalent in M1; dominance more prevalent in CA3.

The BCI task design imposed no constraint on how reward-driving activity should be distributed across the two positive ensemble neurons. In principle, crossing the reward threshold could be supported by the activity of either neuron alone, by both neurons jointly, or by shifting combinations across learning. We therefore characterized each animal’s positive ensemble control strategy by comparing reward-triggered burst probabilities of the two positive ensemble neurons across early and late learning (see Methods). To normalize for regional and learning-dependent differences in activity rates, strategy classification was based on relative rather than absolute burst probabilities. A range of strategies was observed across animals, including dominance of one neuron over the other in driving reward delivery (Figure 2F), cooperation (Figure 2G), dominance-to-cooperation transitions (Figure 2H), and cooperation-to-dominance transitions (Figure 2I). The global distribution of strategies differed between M1 and CA3 (Figure 2J), with CA3 showing a strong bias toward dominance relative to M^52–54^. Within the dominance category, half of all CA3 animals exhibited switching behavior, reversing the identity of the dominant driver between early and late sessions, a pattern absent in M1. M1, by contrast, showed a balance between dominance and cooperation strategies and a greater tendency toward dominance-to-cooperation transitions, in which an initially dominant configuration evolved into a cooperative solution through progressive recruitment of the second positive neuron. No CA3 animal exhibited a purely cooperative solution.

### Learning drives region-specific reorganization of reward-related population activity

While the activity of the direct positive and negative ensemble neurons defines the reward-contingent activity pattern that animals learn to volitionally reactivate, BCI learning induces widespread network reorganization that extends beyond these neurons^22,54–57^. To assess whether M1 and CA3 differ in how learning reshapes activity across the entire recorded network, we compared reward-related tuning in non-ensemble (indirect) neurons (Figure 3A) between the two regions. Reward-modulated neurons were identified from peri-reward calcium PSTHs using a slope-and-threshold criterion that defines bursting probability (Figure 3B; see Methods). A linear decoder trained on the top reward-modulated neurons achieved high classification performance in both regions, whereas decoders trained on non-modulated neurons or temporally shuffled activity performed near chance (Figure 3C), confirming that reward-related information was distributed across the network beyond the positive and negative ensembles, and that the selection criterion isolated neurons carrying reward-timing information through precise temporal patterns rather than elevated firing rates alone.

**Figure 3.**
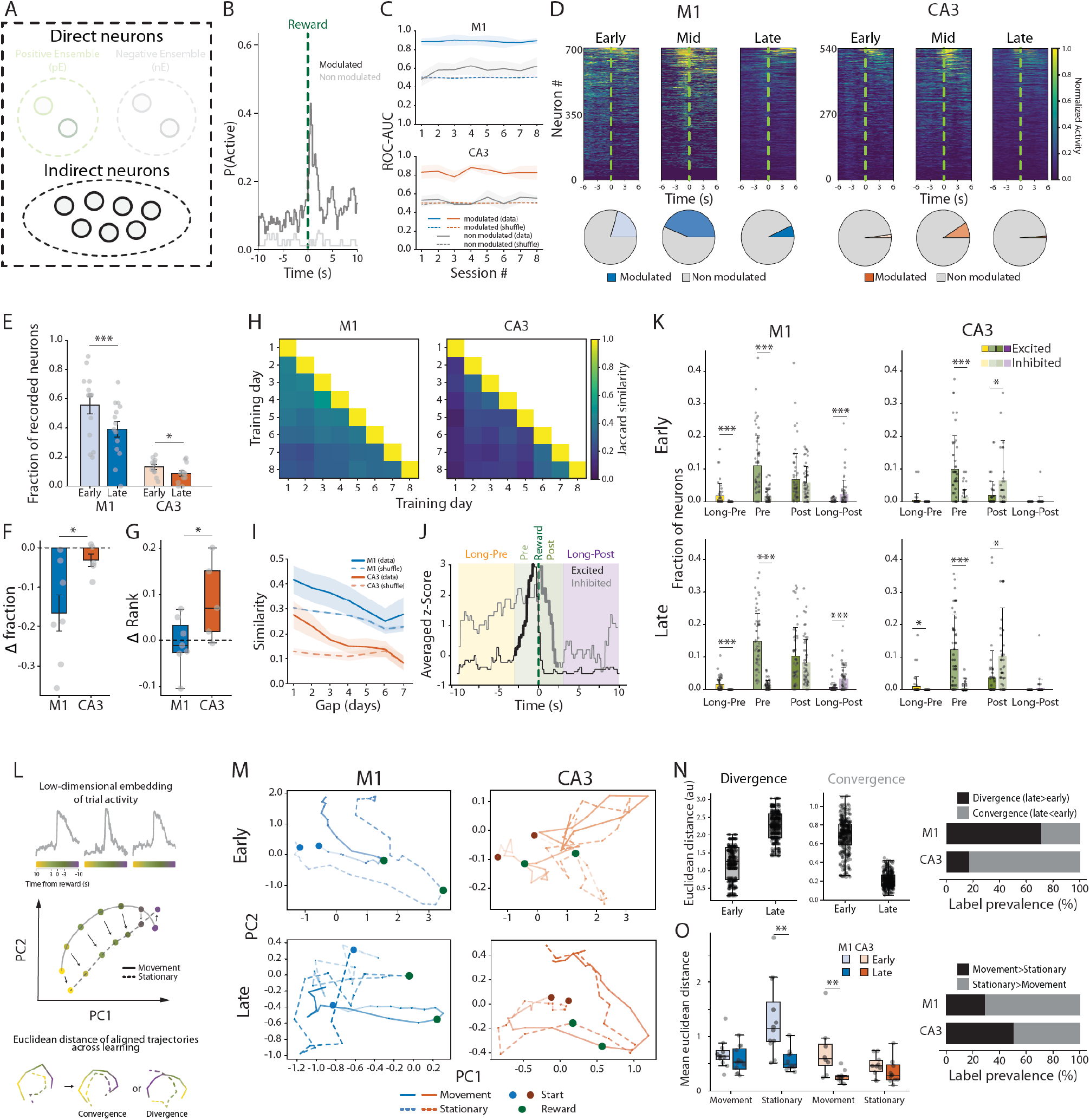
Distinct circuit dynamics reshape reward-related population activity in M1 and CA3 during learning. **A**, Schematic of BCI ensemble architecture, with indirect neurons (i.e. neurons not belonging to neither the positive or negative ensembles) highlighted. **B**, Example peri-reward activity profile for an indirect neuron, showing the probability of being active aligned to reward delivery (t = 0, dashed green line). Dark and light traces, representative reward-modulated and non-modulated neurons, respectively. **C**, Decoding performance (ROC–AUC) of task-modulated neurons across training in M1 and CA3. For each session, decoding evaluated using the top 10 modulated cells and compared with 10 randomly selected non-modulated cells (gray); shuffled controls generated by label permutation. Solid lines, decoding performance from real data; dashed lines, shuffled controls; shading, variability across mice. In both M1 and CA3, decoders trained on the top modulated-cell population showed consistently higher performance than non-modulated or shuffled controls, which remained near chance. **D**, Example reward-aligned modulation profiles across training in M1 and CA3. Heat maps, activity profiles of tracked neurons aligned to reward delivery (green dashed line) during early, mid, and late learning stages in representative M1 and CA3 datasets. Pie charts below each heat map, fraction of neurons classified as modulated on that day. Recording counts: M1 cohort, 30,388 neurons across 7 animals over 54 sessions, with 3,995 unique neurons longitudinally tracked; CA3 cohort, 23,168 neurons across 6 animals over 48 sessions, with 2,773 unique neurons longitudinally tracked. **E**, Fraction of reward-modulated neurons in early and late learning for M1 and CA3. Bars, mean ± s.e.m. across sessions; overlaid points, individual sessions. Group-specific linear mixed-effects models accounting for repeated measurements across mice, with Benjamini–Hochberg correction across groups: significant decreases in the fraction of modulated neurons from early to late learning in both M1 (β = −0.166, FDR-adjusted *p* < 0.001) and CA3 (β = −0.032, FDR-adjusted *p* = 0.035), with a larger reduction in M1. **F**, Change in the fraction of reward-modulated neurons from early to late learning in M1 and CA3, quantified per animal as Δfraction = mean(late) − mean(early). Bars, group means; points, individual animals. Both regions showed a decrease from early to late learning, with a larger reduction in M1. In per-animal analyses, the difference in Delta between regions was significant (paired permutation, p=0.0489). **G**, Change in rank position of modulated cells across learning in M1 and CA3. Δ rank quantifies the change in relative position of modulated cells within recorded network across training. Boxplots, mouse-level Δ rank values; overlaid points, individual mice; dashed horizontal line, no net rank shift. Unpaired permutation test (label shuffling): CA3 exhibited a greater positive rank shift than M1 (*p* = 0.046). **H**, Stability of modulated-cell identity across training days in M1 and CA3. For each mouse, tracked cells were classified as modulated or non-modulated on each training day by thresholding activity, and pairwise similarity in the identity of modulated cells was computed across all day pairs. Heat maps show group-average Jaccard similarity matrices for M1 and CA3, with each entry representing the fraction of modulated cells shared between two training days. Diagonals were set to 1, and only the lower triangle is displayed. Greater persistence of modulated-cell identity across days in M1 than in CA3. **I**, Similarity of modulated-cell ensembles as a function of inter-day interval in M1 and CA3. Solid lines, observed similarity decay across day gaps; dashed lines, shuffled controls (cell identities permuted within each day, preserving the number of modulated cells while destroying cross-day alignment); shading, variability across mice. Permutation test on area under the similarity-versus-gap curve: global group difference significant for Jaccard similarity (*p* = 0.040). Both groups exceeded shuffled-null expectations, consistent with non-random persistence of modulated-cell identity across training days. **J**, Schematic of the epoch-based modulation analysis used to classify reward-related neural responses. Reward-aligned activity partitioned into four epochs relative to reward delivery (green dashed line): Long-Pre (−10 to −3 s), Pre (−3 to 0 s), Post (0 to 3 s), and Long-Post (3 to 10 s). For each cell and epoch, modulation assessed by a directional linear-trend criterion combining slope, correlation, and significance of the fit. Cells classified as excited (significant positive slope and correlation), inhibited (significant negative slope and correlation), or non-modulated. In the example trace, the epoch in which the trend was significant is shown with a thicker line. Counts converted to fractions of total or modulated cells across reward-centered temporal epochs. **K**, Fractions of excited and inhibited neurons across reward-centered temporal epochs in movement rewarded trials, at quarter-session resolution. Classification of excited and inhibited cells as in J. Bars, mean fractions across quarter-session datapoints. Linear mixed-effects models comparing excitatory versus inhibitory fractions, with Benjamini–Hochberg correction across all 16 tests. M1: excited and inhibited fractions differed significantly in Long-Pre, Pre, and Long-Post in both early and late learning (all FDR-adjusted *p* < 0.001), but not in Post (early *p* = 0.434; late *p* = 0.215). CA3: significant differences in Pre in both phases (early and late FDR-adjusted *p* < 0.001), in Post in both phases (early FDR-adjusted *p* = 0.036; late *p* = 0.011), and in Long-Pre during late learning only (FDR-adjusted *p* = 0.045); not significant in Long-Post in either phase (early *p* = 0.332; late *p* = 0.138). **L**, Schematic of the neural-state distance analysis. Population activity from longitudinally tracked neurons projected by PCA into a low-dimensional manifold; trial-aligned trajectories separated by behavioral state (movement versus stationary) and learning stage (early versus late). Neural-state distance, Euclidean separation between paired trajectories at each aligned timepoint in the 4-dimensional PCA space (two dimensions shown for visualization), averaged across timepoints. Changes across learning, defined as late minus early separation within each animal: positive values indicate divergence of movement- and stationary-related trajectories with learning; negative values indicate convergence. **M**, Example neural population trajectories in PCA state space from representative M1 and CA3 mice during early and late learning. Top, early learning; bottom, late learning. Solid lines, movement trajectories; dashed lines, stationary trajectories. M1, blue; CA3, orange. Small filled circles, trajectory start points; green markers, reward-aligned positions. **N**, Left panels, representative learning-related divergence and convergence of movement and stationary neural trajectories. Boxplots show cases where late trajectories became more separated from early ones, classified as divergence (late > early; left example), or closer to early ones, classified as convergence (late < early; right example). Right panel, stacked bar plot summarizing the prevalence of these labels across mice. M1 showed a higher fraction of mice exhibiting divergent trajectories; the CA3 cohort was dominated by convergent trajectories. **O**, Early-versus-late changes in trajectory distance within movement and stationary subspaces in M1 and CA3. Left panel, mean manifold distance for each mouse in M1 movement, M1 stationary, CA3 movement, and CA3 stationary trials. Linear mixed-effects models with phase as a fixed effect, accounting for repeated measurements across mice, with Benjamini–Hochberg correction across the four panel-wise tests. With trials subdivided into two temporal bins, convergence from early to late learning was significant in M1 stationary (β = −0.692, FDR-adjusted *p* = 0.002) and CA3 movement (β = −0.464, FDR-adjusted *p* = 0.002) subspaces, but not in M1 movement (β = −0.117, FDR-adjusted *p* = 0.228) or CA3 stationary (β = −0.047, FDR-adjusted *p* = 0.228) subspaces. Right panel, per-animal prevalence of movement versus stationary trajectory changes, classified by which mean early-to-late shift was larger. M1 showed a higher fraction of mice exhibiting larger movement-related geometric changes; CA3 showed roughly equal proportions of mice with movement- or stationary-related changes.

Learning was associated with a progressive reduction in the fraction of neurons exhibiting significant peri-reward modulation in both M1 and CA3 (Figure 3D-F), indicating that improved task performance coincided with increasingly selective engagement of the recorded network around reward, consistent with previous reports of learning-induced sparsification^4,19,58^. Decoder performance remained stable across all learning sessions despite this reduction (Figure 3C), confirming that sparsification did not degrade the quality of reward-related information carried by the indirect population. The mechanisms underlying sparsification differed between regions. In M1, both reward-modulated and non-modulated neurons showed progressive reductions in peak activity probability, indicating network-wide suppression of activity levels (Extended Data Figure 3A). In CA3, by contrast, non-modulated neurons similarly decreased their peak activity rates, but reward-modulated neurons maintained stable activity levels throughout learning (Extended Data Figure 3A), indicating that CA3 selectively increased the relative difference between the two populations. Consistent with this, the change in activity rank of reward-modulated neurons within the broader recorded population, based on within-session peak activity strength, remained close to zero in M1 but shifted toward significantly higher values in CA3, indicating that modulated neurons came to occupy an increasingly dominant position within the active network in this area (Figure 3G). Representational stability of the reward-modulated population differed markedly between regions, assessed by longitudinal tracking and pairwise Jaccard similarity between binary population vectors across all session pairs (Figure 3H). In M1, cross-session similarity remained substantially above chance across the entire learning period, decaying only gradually with increasing day gap (Figure 3I). In CA3, similarity also exceeded chance at short day gaps but decayed steeply (Figure 3I), indicating that the reward-modulated CA3 population underwent substantially higher rates of representational drift across days than M1^59–62^. Together, these findings indicate that despite arriving at equivalent behavioral outcomes and a shared sparsification, the underlying network-level dynamics differed between regions: population-wide activity suppression and relative representational stability in M1; selective maintenance of modulated neuron activity against a background of network-wide suppression and elevated drift in CA3.

Beyond sparsification, the temporal organization of task-related responses differed markedly between regions. Trials were aligned on reward delivery (t = 0 s), and the trial period was divided into four analysis windows (Long-Pre: −10 s to −3 s; Pre: −3 s to 0 s; Post: 0 s to +3 s; and Long-Post: +3 s to +10 s; see Methods and Figure 3J)^22^. Recorded neurons that showed significant modulation within each window were classified as excited or inhibited based on the sign of their activity change (Figure 3J; see Methods). At the population level, M1 showed a significantly greater proportion of excited neurons in the pre-reward windows (both Long-Pre and Pre), a balanced distribution of excited and inhibited neurons in the Post window, and prevalence of inhibited neurons during Long-Post (Figure 3K). CA3 exhibited a different temporal organization: excited neurons were similarly over-represented in the Pre window, but inhibited neurons were significantly more prevalent in the Post window, with no difference in fraction of modulated neurons at long distances from rewards (Figure 3K). This region-specific temporal fingerprint, with M1 exhibiting sustained pre-reward excitation and CA3 exhibiting a pre-to-post reward excitation-inhibition reversal, was consistent across training (Figure 3K). When trials were split by behavioral state, significant differences emerged selectively across conditions only for M1, which showed greater excitation in movement trials (Extended Data Figure 3B). Contingency degradation experiments comparing threshold-crossing trials without reward and unpredicted reward deliveries without threshold crossings (see Methods) revealed that these peri-reward network dynamics were driven by threshold crossing rather than reward delivery. On threshold-crossing trials without reward, M1 and CA3 neurons maintained temporal modulation patterns consistent with regular BCI performance (Extended Data Figure 3C, top row), whereas unpredicted reward deliveries produced no significant modulation in any temporal window (Extended Data Figure 3C, bottom row). This dissociation was most significant in the post-reward windows, where the characteristic network activity patterns were absent when rewards occurred without threshold crossings, indicating that post-reward network dynamics reflect the consequences of threshold crossings rather than reward processing. Consistently, a decoder trained on threshold-crossing-modulated neurons predicted threshold crossings with high accuracy, significantly exceeding both non-modulated neurons and temporally shuffled controls, but failed to predict reward delivery timing above chance (Extended Data Figure 3D, E), indicating that the region-specific peri-reward dynamics are associated with the threshold-crossing event rather than by reward delivery or learned reward associations.

These circuit-specific peri-reward dynamics were reflected in the geometry of population trajectories across learning. Trial-aligned population trajectories were constructed from peri-reward burst-probability traces of longitudinally tracked reward-modulated neurons, embedded into a single shared principal component analysis (PCA) space per subject across all sessions (Figure 3L, M; trajectories visualized on the first two principal components, analyses computed on the first four: Extended Data Figure 3F, G)^32^. Inter-condition separation, computed as the Euclidean distance between matched timepoints of movement and stationary trial trajectories, calculated separately at early and late learning, increased progressively from early to late sessions in the majority of M1 animals, indicating that the two trajectory types diverged with learning (Figure 3N and Extended Data Figure 3H). By contrast, in the majority of CA3 animals, inter-condition separation decreased with learning, indicating that the two trajectory types converged towards alignment in the state space (Figure 3N and Extended Data Figure 3H). Intra-condition consistency, quantified as the Euclidean distance between matched timepoints of the same trial-type trajectory at early versus late learning, showed opposing patterns across regions. In M1, stationary trials exhibited a significantly greater decrease in intra-condition distance than movement trials, indicating that population activity during stationary trials became substantially more stereotyped with training (Figure 3O). CA3 showed the opposite pattern: movement trials exhibited a significant decrease in intra-condition distance while stationary trials showed no significant change, indicating that movement-associated population activity became substantially more stereotyped with training in CA3 (Figure 3O).

### Equivalent reward-manifold engagement emerges through distinct population dynamics

The circuit-specific temporal organization of peri-reward activity among indirect neurons pointed to distinct exploration dynamics within the reward-contingent region of population state space (the ‘reward manifold’) in M1 and CA3. To track reward manifold access during ongoing behavior, we developed a nonparametric labeling approach in which each population vector at each time bin was assigned a temporal label reflecting its proximity in high-dimensional activity space to peri-reward population states pooled across all reward deliveries, via k-nearest neighbor consensus in high-dimensional activity space (see Methods, Figure 4A and Extended Data Figure 4A, B). This procedure produced a time series of manifold labels, where a label of zero indicates that the current population state resembles the neural configuration observed at reward delivery (“reward state”), and contiguous labeled segments defined reward-manifold traversals (Figure 4B, C and Extended Data Figure 4C). Bins whose nearest neighbors carried no reward-relative label remained unlabeled and defined the intervals separating consecutive traversals. Learning was associated with a marked increase in time spent within the reward manifold in both M1 and CA3: animals generated longer traversals, with a significant increase in total time spent within the reward manifold across sessions (Figure 4D and Extended Data Figure 4D). Reward state crossings also became more frequent across sessions (Figure 4E). With learning, rewarded traversals (traversals during which reward was delivered, Figure 4C) became significantly more frequent and longer than non-rewarded ones in both M1 and CA3 (Figure 4F, G). The conditional probability of receiving a reward during traversals crossing the reward state increased significantly in both regions (Extended Data Figure 4E). Learning also drove the appearance of traversals containing multiple rewards in both M1 and CA3, which were longer than single-reward traversals and appeared predominantly in late sessions (Extended Data Figure 4F).

**Figure 4.**
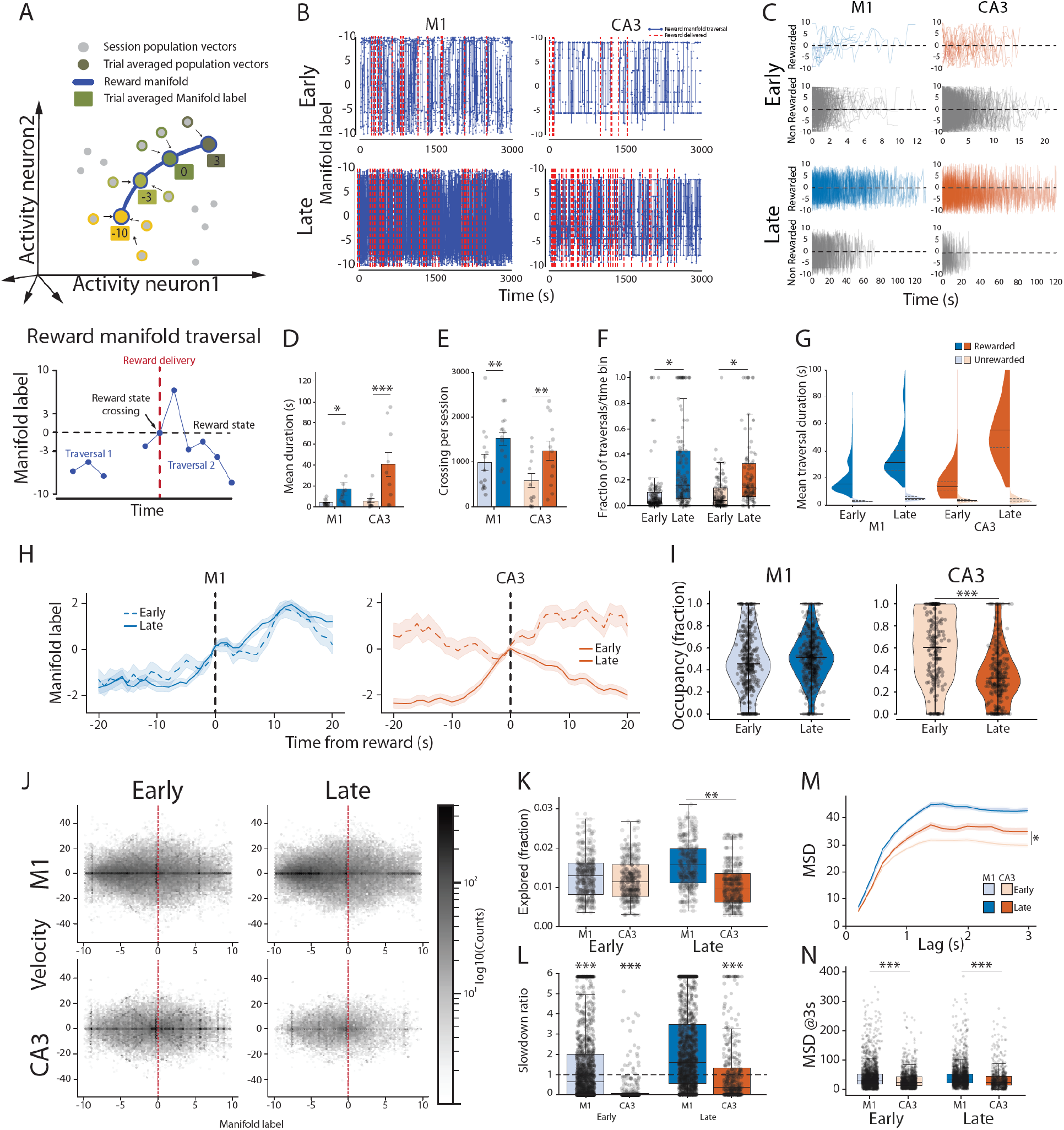
M1 and CA3 exhibit distinct reward-manifold traversal dynamics. **A**, Schematic of reward-manifold identification and traversal analysis. Each population vector at each time bin is assigned a reward-relative label by k-nearest-neighbor consensus in high-dimensional neural state space. The resulting trajectory of reward-relative labels defines manifold traversals as contiguous labeled segments; unlabeled bins, during which population activity cannot be associated with the reward manifold, separate consecutive traversals. The schematic illustrates two distinct traversals separated by an unlabeled pause. **B**, Example session-wise reward-manifold state labels in early and late learning for M1 and CA3. Blue traces, timepoints assigned a reward-relative manifold label (procedure in A), plotted across representative sessions. y axis, reward-relative manifold position label; 0, reward-centered state boundary. Red dashed lines, reward delivery times. Top, early learning; bottom, late learning. Late sessions in both regions showed denser occupancy of labeled reward-manifold states across the session. **C**, Example rewarded and non-rewarded manifold traversals in the first and last training sessions for M1 and CA3. Traversals aligned to onset and grouped by whether they contained reward delivery (rewarded) or not (non-rewarded). Left, first session; right, last session. Top two rows, M1; bottom two rows, CA3. Dashed horizontal lines, reward-centered state boundary. In both regions, rewarded traversals in late sessions were more prolonged and more densely represented than in early sessions, whereas non-rewarded traversals remained shorter and less sustained. **D**, Mean duration of individual reward-manifold traversals in early and late learning for M1 and CA3 (segments >300 s excluded). Bars, mean ± s.e.m. across sessions; overlaid points, individual sessions. Linear mixed-effects models accounting for repeated measurements across mice: significant increases in mean duration from early to late learning in both M1 (β = 13.5 s, FDR-adjusted *p* = 0.021) and CA3 (β = 33.6 s, FDR-adjusted *p* < 0.001). **E**, Number of reward-manifold zero crossings (sign changes across the reward-centered boundary) per session in early and late learning for M1 and CA3. Bars, mean ± s.e.m. across sessions; overlaid points, individual sessions. Linear mixed-effects models accounting for repeated measurements across mice: significant increases in zero crossings from early to late learning in both M1 (β = 543, FDR-adjusted *p* = 0.005) and CA3 (β = 669, FDR-adjusted *p* = 0.005). **F**, Fraction of rewarded manifold traversals in early and late learning for M1 and CA3, quantified in 5-min bins. Boxplots, bin-level values; overlaid points, individual bins. Two-sided permutation tests with within-mouse label swapping, with Benjamini–Hochberg correction across groups: fraction of rewarded traversals increased from early to late learning in both M1 (early mean = 0.100, late mean = 0.306, FDR-adjusted *p* = 0.046) and CA3 (early mean = 0.091, late mean = 0.241, FDR-adjusted *p* = 0.046). **G**, Duration of rewarded and non-rewarded reward-manifold traversals across learning in M1 and CA3. Individual traversal durations (≤300 s) classified by reward status (rewarded vs non-rewarded) and learning phase (early vs late); violin plots. Mixed-effects models testing planned contrasts, accounting for repeated measurements across mice and sessions, with Benjamini–Hochberg correction across 12 comparisons. Rewarded traversals were significantly longer than non-rewarded ones in both groups and both phases (M1 early, M1 late, CA3 early, CA3 late: all FDR-adjusted *p* < 0.001). Significant early-to-late increase in traversal duration was detected only for rewarded traversals in M1 (β = 28.8 s, FDR-adjusted *p* = 0.033); no significant early-to-late change for M1 non-rewarded traversals, CA3 rewarded traversals, or CA3 non-rewarded traversals. No significant differences between M1 and CA3 within matched phase and reward-status conditions after correction. **H**, Event-triggered averages of continuous reward-manifold state trajectories in M1 and CA3 across learning. Reward-state and rewarded population-state traces aligned to transitions across the reward state, averaged across events in early and late learning. Dashed traces, early learning; solid traces, late learning; shading, variability across events. Vertical black lines, aligned reward time. **I**, Post-reward occupancy of positive reward-manifold states within rewarded traversals in early and late learning for M1 and CA3. Violin plots, distribution of post-reward occupancy values; overlaid points, individual traversals. Linear mixed-effects models fit separately for M1 and CA3, accounting for repeated measurements across mice and sessions, with Benjamini–Hochberg correction across groups. In M1, post-reward positive occupancy did not change significantly from early to late learning (early mean = 0.477, late mean = 0.506, β = 0.027, FDR-adjusted *p* = 0.532). In CA3, post-reward positive occupancy decreased significantly from early to late learning (early mean = 0.587, late mean = 0.353, β = −0.251, FDR-adjusted *p* < 0.001). **J, J**oint state-position and velocity occupancy during rewarded return traversals in M1 and CA3 across learning. Log-density hexbin maps of midpoint state-position and instantaneous velocity samples from rewarded return traversals, plotted on a common axis range and binning grid for four conditions (M1 early, M1 late, CA3 early, CA3 late). Dashed vertical line, x = 0 (reward-state boundary). Across learning stages, CA3 occupancy remained more tightly concentrated within a narrower range of positions and velocities, consistent with spatially confined dynamics; M1 occupied a broader region of the position–velocity plane and spanned wider velocity values, consistent with less confined, more continuously traversing dynamics. **K**, Explored position–velocity state space for rewarded traversals in M1 and CA3 across learning. Boxplots, traversal-level explored state-space fraction (proportion of grid bins visited by each traversal on a shared position–velocity grid); overlaid points, individual traversals. Early in learning, M1 and CA3 showed similar explored-space fractions (M1 *n* = 234 traversals, CA3 *n* = 138 traversals; mouse-cluster permutation test on median difference, *p* = 0.741). By late learning, M1 traversals explored a significantly larger fraction of the joint state space than CA3 (M1 *n* = 278, CA3 *n* = 96; median difference = 0.005, *p* = 0.006, FDR-adjusted *p* = 0.012). **L**, Per-traversal slowdown around the state boundary in M1 and CA3 during early and late learning. Slowdown ratio = median speed near the boundary (lowest 20% of |x| values) divided by median speed far from the boundary (60–90% range of |x|); values below 1 indicate boundary-centered slowing. Boxplots, traversal-wise slowdown ratios; overlaid points, individual traversals; dashed line, ratio of 1. Within-condition near-versus-far speed comparisons showed significant slowdown in M1 early (*n* = 1,319, Wilcoxon signed-rank *p* < 0.001), CA3 early (*n* = 548, *p* < 0.001), and CA3 late (*n* = 449, *p* < 0.001), but not in M1 late (*n* = 1,525, *p* = 1.00). Between-region comparisons (M1 vs CA3, Mann–Whitney U): early, *n*_M1 = 1,220, *n*_CA3 = 522, median ratio 0.655 vs 0.000, *p* < 0.001; late, *n*_M1 = 1,405, *n*_CA3 = 398, median ratio 1.59 vs 0.367, *p* < 0.001. **M**, Mean-squared displacement (MSD) of rewarded traversals in M1 and CA3 across learning, computed per traversal up to a lag of 3.0 s. Mean MSD curves shown for early and late learning. Global whole-curve permutation test (L2 distance between early and late mean curves under within-mouse label shuffling): M1 showed no significant early-to-late change in MSD shape (*p* = 0.746); CA3 showed a significant global curve shift across learning (*p* = 0.023). **N**, Traversal-wise MSD at 3 s, a scalar summary of short-timescale displacement magnitude. M1 exhibited larger MSD at 3 s than CA3 in both early (median: M1 = 31.4, CA3 = 22.8; Mann–Whitney U, *p* < 0.001) and late learning (median: M1 = 33.1, CA3 = 24.9; *p* < 0.001).

Aligning manifold label trajectories to reward-state crossing events revealed a learning-dependent dissociation between regions (Figure 4H). In M1, manifold labels increased monotonically from negative to positive values across the reward-aligned window regardless of learning stage, closely approximating a diagonal and indicating that M1 population states traversed the reward manifold in strict chronological order, with the reward state representing a waypoint in a continuous directed flow. In CA3, this relationship was learning-dependent: during early learning, population states hovered near the reward state both before and after reward delivery without a clear temporal structure; during late learning, pre-reward states exhibited negative labels that rose through the reward state before declining, describing a non-monotonic approach-and-return dynamic. Consistent with this divergence in post-crossing dynamics, positive-side occupancy differed between regions (Figure 4I). In early learning, positive-side occupancy following reward state crossing was comparable across M1 and CA3. With learning, M1 positive-side occupancy remained stable, while CA3 positive-side occupancy decreased significantly.

Whether the approach-and-return dynamic in CA3 reflected genuine looping motion or merely temporal-label asymmetry was tested by analyzing trajectory kinematics on the reward manifold. Genuine looping motion predicts a distinctive kinematic signature: trajectories should slow as they approach the turning point near the reward state and accelerate as they leave, producing local deceleration; through-passage trajectories should instead maintain speed across the same region. Joint position-velocity density maps pooled across traversals confirmed the predicted CA3 signature: activity concentrated near the reward state position at low velocity in both early and late sessions, whereas M1 velocity was distributed broadly across the manifold regardless of learning stage (Figure 4J). With learning, M1 traversals sampled a substantially larger region of joint position-velocity space than CA3 (Figure 4K). Slowdown ratios at the reward state were consistently below or equal to 1 in CA3 throughout training, indicating deceleration near the reward state; M1 traversals instead exhibited ratios around 1 by the end of learning, indicating no slowdown during crossing of the reward state (Figure 4L). Mean squared displacement (MSD) of rewarded traversals was greater in M1 than in CA3 at both early and late learning (Figure 4M, N), indicating that M1 population activity explored a substantially larger extent of the reward manifold on a traversal-by-traversal basis. The MSD curve was stable across learning in M1 but increased significantly in CA3, indicating that learning progressively expanded the spatial footprint of CA3 traversals within the reward manifold.

### Circuit architecture shapes the dynamical implementation of learning

The convergence of equivalent behavioral outcomes from circuit-specific population dynamics raises the question of whether circuit architecture alone, rather than task structure, is sufficient to determine the population-level implementation of volitional learning. We tested this by implementing a minimal closed-loop BCI task *in silico*, training distinct recurrent neural networks (RNNs) under minimal constraints chosen to instantiate high-level differences in recurrent circuit computational structure and dynamical regime between M1 and CA3: a low-rank recurrent structure for the M1-like model, capturing low-dimensional trajectory generation^31,32,34,36,63–65^, and a Hopfield-like symmetric autoassociative structure for the CA3-like model, capturing multiple point attractors and pattern completion at retrieval (Figure 5A and Extended Data Figure 5A-G)^26,37,39,41,66^. Population-geometry metrics were quantified using analysis readouts matched to those carried out on the biological data.

**Figure 5.**
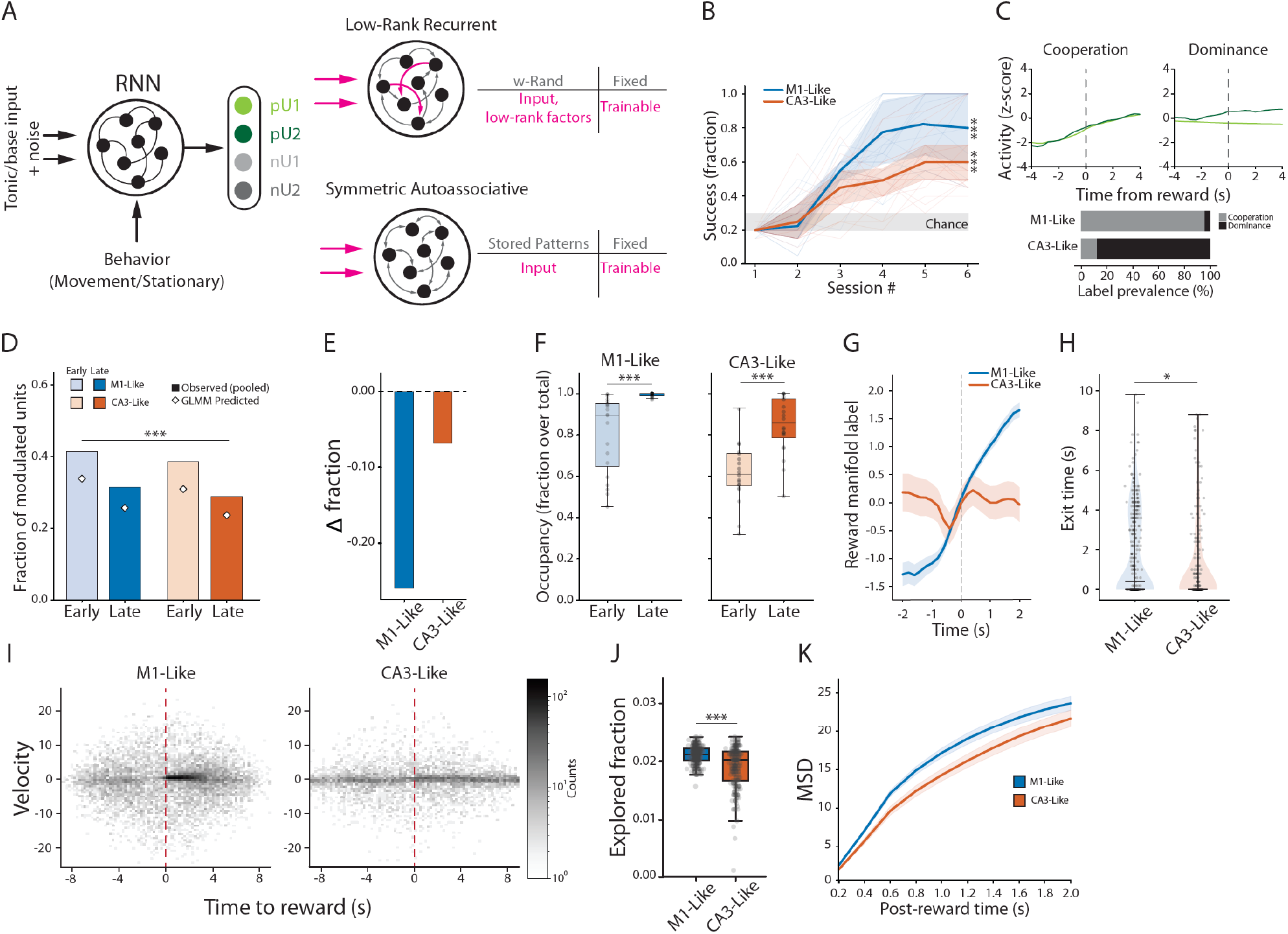
Minimal recurrent architectures reproduce distinct learning dynamics *in silico*. **A**, Schematic of the task and model comparison. Both networks received the same tonic/base input, behavior-dependent context (movement versus stationary), and noise, and were read out through a fixed four-unit ensemble with two positively weighted (pU1, pU2; shades of green) and two negatively weighted (nU1, nU2; shades of grey) units. The M1-like model used a low-rank recurrent perturbation on a fixed random backbone, with input and low-rank factors trainable; the CA3-like model used a fixed symmetric autoassociative recurrent core built from stored patterns, with input pathways trainable. **B**, Learning curves across six sessions for 20 seeds per model. Thin lines, individual seeds; thick lines, median across seeds; shaded bands, interquartile range. Both models started from the same baseline reward fraction, corresponding to chance performance (0.2), and improved with training. Mean success fraction across the final two sessions tested against chance (one-sided Wilcoxon signed-rank test, Bonferroni-corrected): both models performed significantly above chance at the end of learning (M1-like models, *n* = 20, *p* < 0.001; CA3-like models, *n* = 20, *p* < 0.001). **C**, Reward-aligned positive-readout-unit dynamics and late control regime. Top, representative reward-aligned traces illustrating two distinct late regimes: cooperative recruitment of the two positive readout units, versus dominant recruitment of a single positive unit. Bottom, prevalence of these late labels across seeds. M1-like models, driving was cooperative (95.0% co-driving, 5.0% single-driving); CA3-like models were dominated by single-unit drive (88.6% single-driving, 0% co-driving, 11.4% other). **D**, Fraction of modulated units in early and late phases of training for the M1-like and CA3-like models, with neuron-level generalized linear mixed-model predictions overlaid. Mixed-model inference: significant main effects of model (*p* < 0.001) and phase (*p* < 0.001); no model × phase interaction (*p* = 0.640). **E**, Late-minus-early change in modulation prevalence in the M1-like and CA3-like models. Median delta from pooled fractions: M1-like = −0.262; CA3-like = −0.070. Larger reduction in M1-like. **F**, Reward-manifold occupancy in early and late learning for the M1-like and CA3-like models. Occupancy, fraction of non-NaN reward-manifold labels. Early-to-late occupancy increased significantly in both models (M1-like: paired *t*-test *p* < 0.001; Wilcoxon *p* < 0.001; CA3-like: paired *t*-test *p* < 0.001; Wilcoxon *p* < 0.001). **G**, Reward manifold dynamics in M1-like and CA3-like models, restricted to rewarded trials whose post-reward manifold-label trajectory crossed below 0 at least once within +10 s (matching the rewarded-returning subset used in the biological data). Solid lines, mean trajectory. **H**, Rewarded-returning exit-time distribution (exit time = first post-reward time at which the manifold label fell below 0). M1-like models exhibited longer late rewarded-returning exit times than CA3-like (event-level Mann–Whitney *p* = 0.004; *n* = 311 returning events in M1-like, 213 in CA3-like; seed-mean comparison *p* = 0.018). **I**, Joint position–velocity occupancy maps for rewarded return traversals in M1-like and CA3-like models during late learning. Log-scaled grayscale; darker, higher sample density. Vertical dashed line, reward-state boundary. M1-like trajectories occupied a broader region of joint state space than CA3-like, whereas CA3-like occupancy remained more concentrated around lower velocities and a narrower range of manifold positions. **J**, Traversal-level explored state-space fraction (fraction of visited bins on the common position–velocity grid from I) for rewarded-returning traversals in M1-like and CA3-like models. M1-like traversals explored a larger fraction of joint state space than CA3-like (*p* < 0.001). **K**, Mean-squared displacement (MSD) of rewarded-returning post-reward trajectories over the first 2 s in M1-like and CA3-like models. After learning, M1-like models exhibited larger short-lag MSD than CA3-like (AUC comparison *p* = 0.017; median AUC difference M1 − CA3 = 5.41; Cliff’s δ = 0.445), with FDR-significant lag-wise differences from 0.2 to 1.6 s.

Across measured readouts, the models reproduced key signatures of the biological dissociation. Both models learned to obtain rewards through modulation of a fixed readout ensemble (Figure 5B). The strategy used to drive the readout units dissociated between models: the M1-like network recruited readout units in a distributed cooperation regime, while the CA3-like network concentrated drive onto a single unit in a dominance regime (Figure 5C). Beyond the readout units, population activity sparsified with learning in both architectures (Figure 5D, E). Reward manifold occupancy increased from early to late sessions in both models (Figure 5F), paralleling the in vivo result. Reward manifold traversal dynamics dissociated between models: the M1-like network showed sustained and amplified positive manifold occupancy following threshold crossing, with crossing itself embedded in a monotonically increasing sequence from negative to positive manifold labels and the reward state as a waypoint in this trajectory (Figure 5G). The CA3-like network instead hovered near baseline, producing a briefer positive traversal upon threshold crossing with earlier return toward pre-reward configurations (Figure 5G). Consistently, post-crossing occupancy of the positive manifold portion was significantly longer in the M1-like than the CA3-like network, reproducing the biological asymmetry (Figure 5H). The M1-like model traversals sampled a substantially larger region of joint position-velocity space than the CA3-like model ones (Figure 5I, J). Mean squared displacement analysis of the reward manifold coordinate paralleled the biological result: activity in the CA3-like model explored a substantially smaller reward manifold extent than the M1-like model in the early aftermath of reward delivery, a difference that became non-significant at longer lags (Figure 5K). Overall readouts such as learning performance, readout ensemble control, exploration of the reward manifold, and post-reward occupancy of the positive manifold space were robust to network sizes and activation function selection (Extended Data Figure 6).

## Discussion

Our results reveal that distinct circuits learn the same volitional associative learning task by arriving at equivalent outcomes with fundamentally different dynamical implementations. In both M1 and CA3, learning was accompanied by a set of shared network signatures (modulation of reward-controlling neurons, network-level sparsification, and greater exploration of reward-related activity patterns), each underpinned by mechanistically distinct implementations (distributed cooperation versus dominance in ensemble control, geometric divergence versus convergence of behavioral-state trajectories, and continuous trajectory flow versus approach-and-return dynamics on the reward manifold). Recurrent neural network models endowed with distinct minimal constraints that recapitulate the dominant computational regime of each region, and trained on an *in silico* BCI task, captured key features of the shared and divergent solutions observed in biological networks, suggesting that local circuit architecture constrains the form of the observed dynamical dissociation. These results are consistent with the interpretation that the opposing dynamical signatures observed during learning reflect the native computational architectures of each region: M1 operating as a trajectory-generating low-dimensional dynamical system in which population activity flows through sequential states^31,32,36,67^, and CA3 as an autoassociative network in which population dynamics converge toward stable configurations^26,37,39,41,66^. Learning thus recruits each circuit’s native dynamical regime in service of the same learning goals through distinct population dynamics and geometries, consistent with principled degeneracy in neural systems^13–18,68^.

In both regions, animals achieved successful resolution of the credit assignment problem inherent in ensemble BCI control (identifying which neural configuration produces reward and learning to reliably recruit it)^5,23,51^ through mechanisms consistent with their respective circuit architectures. In M1, the tendency toward progressively equivalent recruitment of both positive ensemble neurons into co-driving configurations^19^ reveals a flexible redistribution across the neurons involved, a solution consistent with a distributed population coding. The discrete commitment to a single driver and the pattern of switching between early and late learning indicate that CA3’s control strategy is consistent with convergence onto dominant configurations and winner-take-all transitions^26,39,69,70^. Similarly, equivalent sparsification of reward-related network activity emerged through mechanistically distinct routes. In M1, proportional suppression of both modulated and non-modulated neurons is consistent with a global modulation of excitability^22,56^. The selective stabilization of reward-related activity against broad network suppression in CA3 is consistent with recurrent amplification of task-relevant signals^59^, a feature of autoassociative organization.

The temporal organization of peri-reward responses reflected each circuit’s dynamical regime. In M1, sustained pre-reward excitation transitioned to a post-reward balance between excitation and inhibition, consistent with a trajectory-generating system that continues to flow through state space rather than actively suppressing activity after reward^31^. In CA3, pre-reward excitation was followed by active post-reward inhibition, consistent with convergence onto a reward-associated state followed by active release^41,70^. Despite near-complete turnover of the reward-modulated CA3 population across sessions, the temporal structure of peri-reward responses was consistent across the learning period sampled, as it was in M1, where representational stability^55,61^ accompanied a similarly consistent temporal fingerprint. The persistence of these fingerprints across changes in cellular identity indicates that they reflect circuit-level dynamical organization rather than representational structures progressively shaped by learning.

Learning also reorganized the geometry of population activity. In M1, progressive differentiation of movement- and stationary-trial trajectories indicates that M1 incorporates behavioral state into its trajectory-generating dynamics, producing an increasingly behavior-sensitive population geometry^25,71^. In CA3, the reduction in trajectory separation with learning suggests that CA3 develops more behaviorally invariant task representation, consistent with an autoassociative network that abstracts over locomotion to retrieve a stable configuration^69,70^. Together with the temporal observations, these opposing reorganizations indicate that M1 incorporates behavioral context into its dynamics during volitional learning, whereas CA3 abstracts away from it to converge onto stereotyped population configurations.

The dynamics of reward manifold traversal reveal the clearest expression of each circuit’s computational identity during task performance. In M1, the embedding of the reward state within a continuous directed flow is consistent with a system that generates sequential trajectories through state space that map trial progression^31,32,34,36^. In CA3, the approach-and-return excursion is consistent with a system that performs discrete state transitions rather than continuous flow to map task structure^3,41,70,72^. The correspondence between this looping motion and the temporal profile of the control ensemble’s state (Extended Data Figure 2D) suggests that CA3’s autoassociative architecture might propagate the direct neuron’s temporal structure across the broader recorded network as a coherent, network-wide signal^37,41,59^. Recurrent network models instantiating high-level computational differences between the two regions captured key features of the population-level dissociation observed in vivo. Both M1-Like and CA3-Like architectures arrived at the shared biological endpoints (reward acquisition, population sparsification, and reward manifold engagement) while diverging in ensemble control strategy and reward manifold traversal dynamics in ways that paralleled the biological dissociation: distributed co-driving and continuous trajectory flow with the reward state as a waypoint, versus single-unit dominance and approach-and-return dynamics with restricted state-space exploration following threshold crossing, respectively. That these dissociations emerge from networks differing in recurrent connectivity structure and dynamical regime indicates that architectural differences are sufficient to constrain the population-level form of learned solutions^68,73^.

Together, these findings indicate that associative learning is achieved through circuit-specific dynamical implementations shaped by architectural constraints. Shared features across circuits reflect the invariant requirements of associative learning, whereas divergent features reflect dynamical constraints associated with each circuit’s organization. These results support the idea that principled degeneracy, in which architecturally distinct circuits arrive at equivalent learning outcomes through fundamentally different population-level mechanisms, represents a feature of learning in both biological and artificial neural systems.

## Acknowledgements

We thank the animal facility, the research instrumentation facility, and the mechanical and electrical workshops of the Biozentrum of the University of Basel for excellent technical assistance. We thank the whole Donato lab for the productive discussions and feedback on the project. We thank Liset de la Prida, Everton Joao Agnes, Mathias Mann, and Albert Tsao for their insightful feedback on the manuscript. A.d.V. and F.D. are grateful to Andreas Luthi and Mathias Mann for their support during the completion of this work.

## Author contributions

F.D., C.M., and A.d.V. conceived the study. C.M. wrote and optimized the Open-CaBCI software. A.d.V., C.M., R.V.M., and L.M. acquired data. A.d.V., C.M., and F.D. analysed the data. A.d.V., C.M., and F.D. prepared figures. M.C-R. helped with the initial development of the BCI task. D.B. helped with the kinematic analysis. G.B. helped with the RNN model development and refinement. F.D. wrote the original draft of the manuscript, with help from A.d.V. and C.M., and feedback from all authors.

## Data and materials availability

All data will be made available in the manuscript, the supplementary material, or a publicly accessible database.

---

**Extended Data Figure 1.**
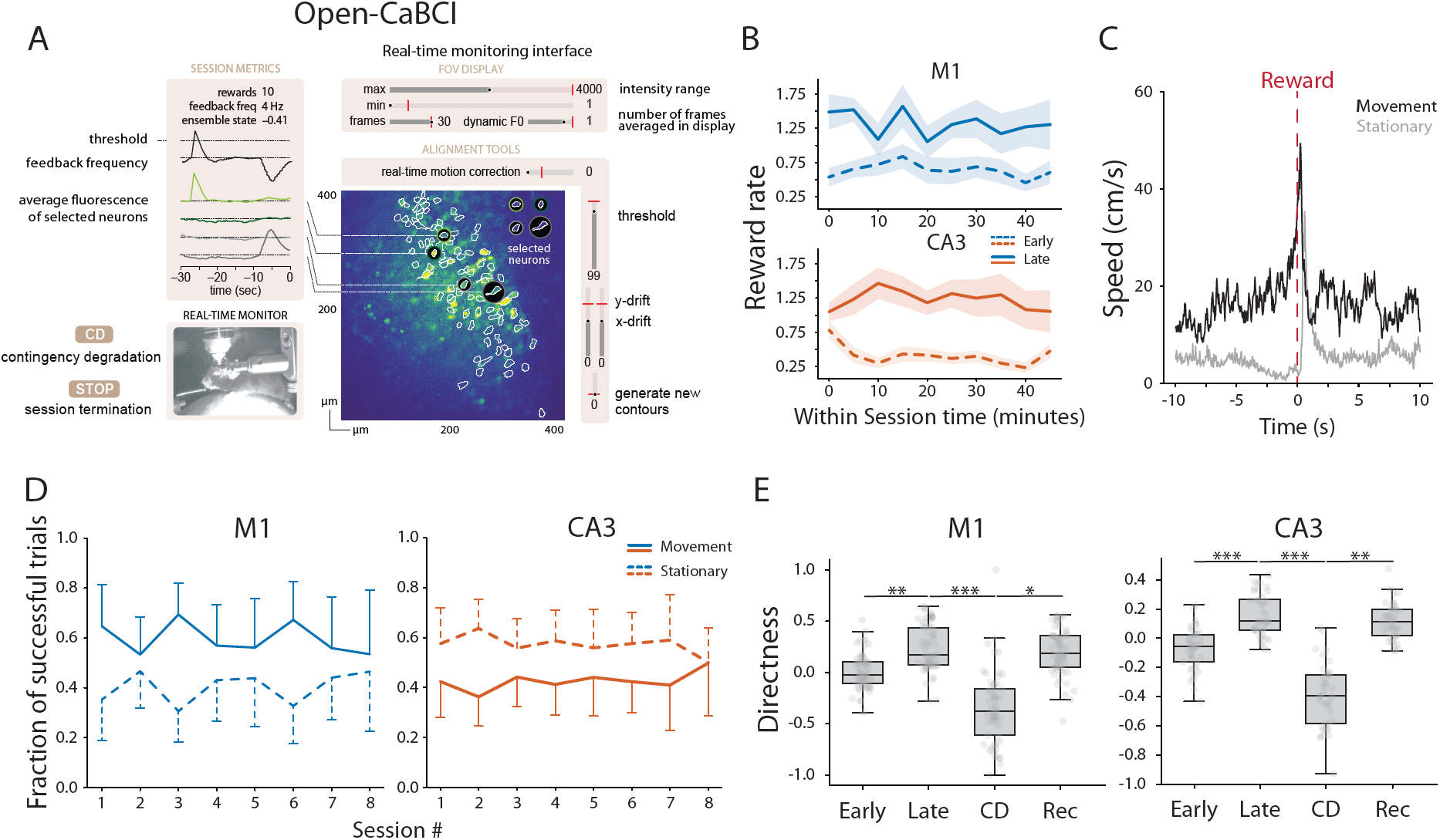
BCI control is independent from locomotion and associated with activity-reward contingency. **A**, Real-time graphical user interface from Open-CaBCI, used to monitor and control task performance during BCI sessions. The interface displayed online session metrics, including reward count, auditory feedback frequency, ensemble-state value relative to threshold, and the fluorescence activity of selected neurons. The field-of-view panel visualized identified neuronal contours and highlighted the subset of neurons used for online control, together with tools for adjusting display parameters, monitoring motion correction, tracking x–y drift, setting reward threshold, and updating contours. A live behavioral camera feed allowed continuous monitoring of the animal, and additional controls enabled contingency-degradation manipulation and manual session termination. **B**, Within-session reward dynamics during early and late phases of BCI learning in M1 (top) and CA3 (bottom). Reward rate computed in consecutive 5-min bins within each session and averaged across the first two training sessions (early) and the last two training sessions (late). Solid lines, late sessions; dashed lines, early sessions; shading, mean ± s.e.m. across animals. Mixed-effects model accounting for repeated measurements across mice and sessions: significant increase in reward rate from early to late (β = 0.838, *p* < 0.001); no significant phase × group interaction (β = −0.160, *p* = 0.453). Within-session time interactions also not significant (phase × time, *p* = 0.262; group × phase × time, *p* = 0.280). **C**, Example locomotor-speed profiles aligned to rewarded-trial onset (time 0; dashed line), from 10 s before to 10 s after, for trials preceded by locomotion (black, nivement) or immobility (gray, stationary). **D**, Fraction of successful movement or stationary trials across training sessions in M1 (left) and CA3 (right). Solid lines, movement-preceded trials; dashed lines, stationary-preceded trials; error bars, mean ± s.d. across animals. Mixed-effects model on chunk-level (5-min) data, accounting for repeated measurements across mice and sessions: no effect of training phase on the fraction of movement-preceded successful trials (β = 0.055, *p* = 0.620), no main effect of group (β = 0.200, *p* = 0.271), and no group × phase interaction (β = −0.075, *p* = 0.617). Group-specific late-versus-early contrasts: M1 β = −0.017, *p* = 0.881; CA3 β = 0.058, *p* = 0.535. **E**, Directness of auditory-feedback trajectories across learning (Early, Late), contingency degradation (CD), and recovery (Rec) phases in M1 (left) and CA3 (right). Directness as in Figure 1G. Boxplots, bin-level directness values; overlaid points, individual bins. Linear mixed-effects models performed separately in M1 and CA3, accounting for repeated measurements across mice and sessions: directness increased from Early to Late (M1 *p* = 0.004; CA3 *p* < 0.001), decreased during CD (M1 *p* < 0.001; CA3 *p* < 0.001), and recovered in Rec sessions (M1 *p* = 0.015; CA3 *p* = 0.002). Benjamini–Hochberg-corrected pairwise comparisons: significant differences between Late vs CD and between CD vs Rec in both groups; Late vs Rec not significant, indicating disruption of trajectory efficiency during CD and recovery toward late-learning levels.

**Extended Data Figure 2.**
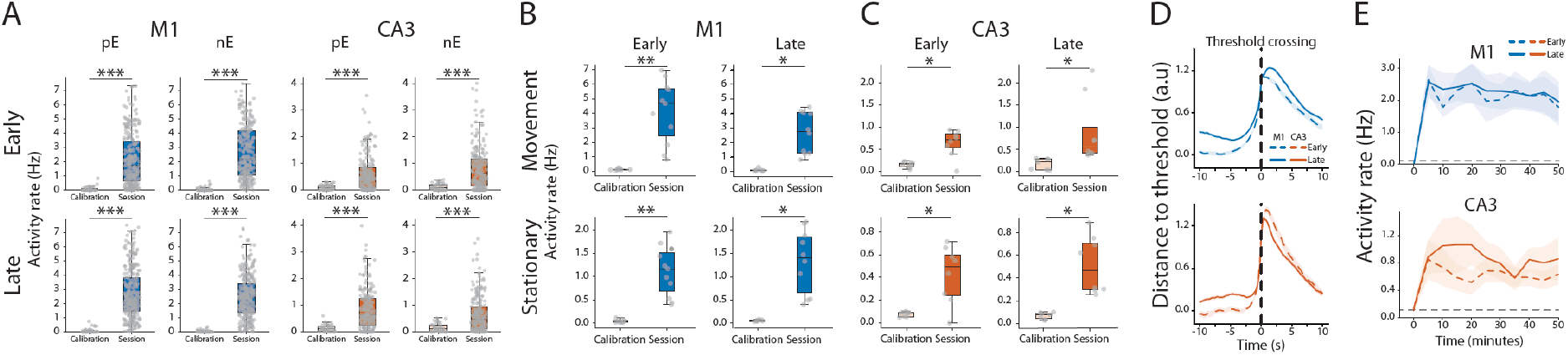
Task-specific ensemble recruitment underlies BCI control in both M1 and CA3. **A**, Firing rates of positive and negative ensemble neurons during calibration and BCI-session epochs in M1 and CA3. Panels organized by region, phase, and ensemble class; boxplots, 5-min bin values with overlaid datapoints. Linear mixed-effects models with context (calibration versus session) as a fixed effect, accounting for repeated measurements across animals, with Benjamini–Hochberg correction across the eight panel-wise tests. Session bins showed significantly higher firing rates than calibration bins in every subgroup: M1 early positive (β = 1.50 Hz, FDR-adjusted *p* < 0.001), M1 early negative (β = 2.16 Hz, FDR-adjusted *p* < 0.001), M1 late positive (β = 2.16 Hz, FDR-adjusted *p* < 0.001), M1 late negative (β = 2.19 Hz, FDR-adjusted *p* < 0.001), CA3 early positive (β = 0.510 Hz, FDR-adjusted *p* < 0.001), CA3 early negative (β = 0.704 Hz, FDR-adjusted *p* < 0.001), CA3 late positive (β = 0.695 Hz, FDR-adjusted *p* < 0.001), CA3 late negative (β = 0.517 Hz, FDR-adjusted *p* < 0.001). For calibration periods, no significant positive-versus-negative differences were detected after correction (M1 early β = 0.012, FDR-adjusted *p* = 0.695; M1 late β = 0.037, FDR-adjusted *p* = 0.430; CA3 early β = −0.011, FDR-adjusted *p* = 0.695; CA3 late β = −0.020, FDR-adjusted *p* = 0.695). **B**, Activity rates of ensemble-positive neurons during movement-preceded (top) and stationary-preceded (bottom) trials in M1 during calibration and BCI-session epochs. Light shading, calibration; dark shading, training session. Left, early stage; right, late stage. Boxplots, session-level distributions with overlaid points indicating individual sessions. Paired sign-flip permutation tests matched by animal and session: activity rates significantly higher during the session than during calibration for both movement- and stationary-preceded trials in early learning (*p* = 0.008 for both) and late learning (*p* = 0.015 for both). **C**, Same as B, but for CA3. Activity rates significantly higher during the session than during calibration for both movement- and stationary-preceded trials in early learning (*p* = 0.030 for both) and late learning (*p* = 0.015 for both). **D**, Ensemble state signal normalized to session reward threshold (dimensionless distance to threshold), aligned to reward delivery (vertical dashed line, time 0). Peri-reward window: −10 to +10 s. Lines, mean ± s.d. across reward events pooled by region and learning phase. Dashed lines, early learning; solid lines, late learning. Both M1 and CA3 showed a sharp rise toward threshold before reward, followed by post-crossing relaxation away from threshold. **E**, Activity rate of positive ensemble neurons across the full session in M1 (top) and CA3 (bottom). 5-min bins from calibration through the BCI session. Solid lines, mean activity rate across late sessions; dashed lines, early sessions; shading, variability across sessions. In both regions, ensemble-positive activity increased sharply after session onset and remained elevated throughout the session.

**Extended Data Figure 3.**
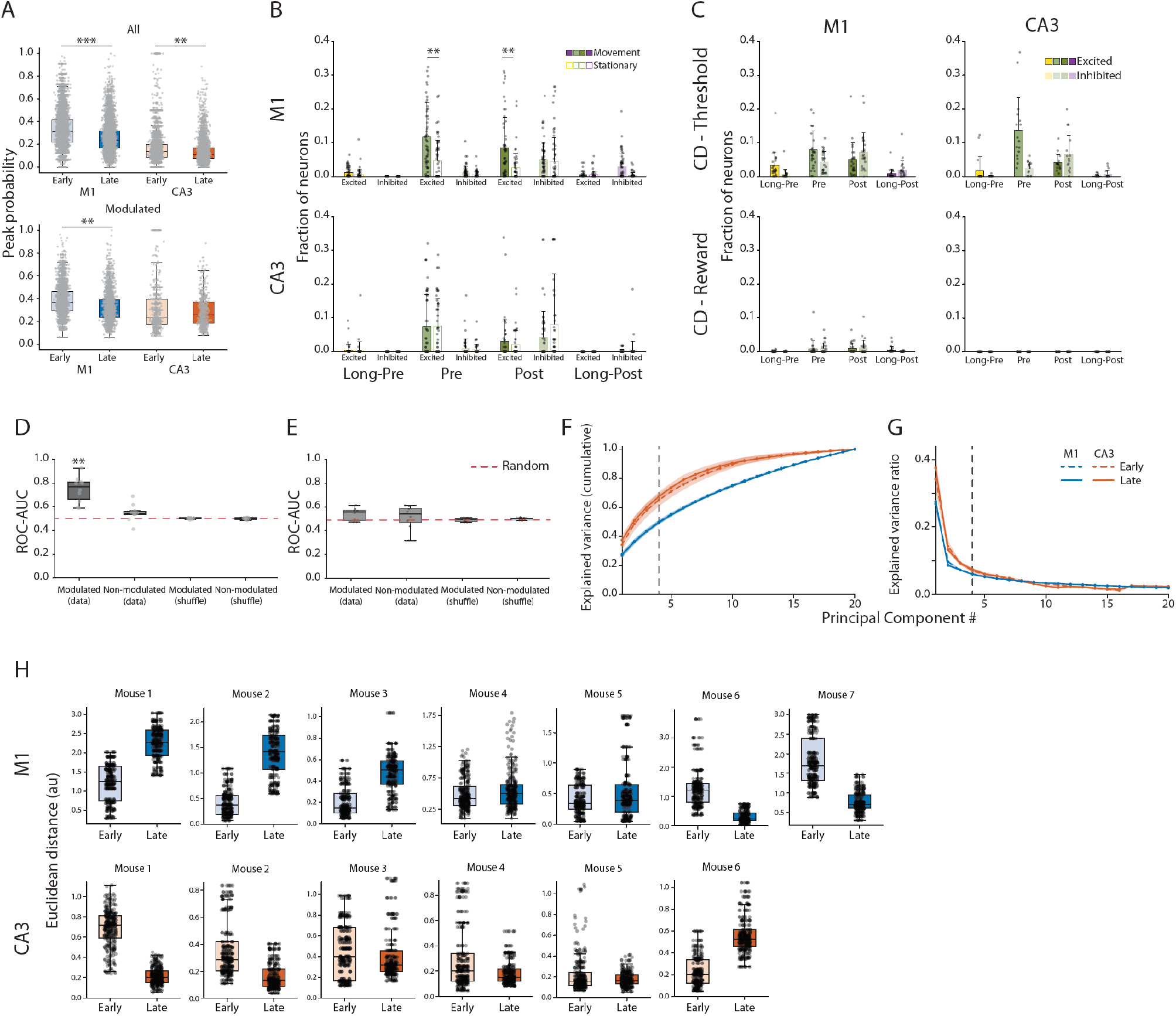
Shared reward-related network reorganization emerges through distinct circuit mechanisms. **A**, Peak response probability of reward-aligned activity in all cells and task-modulated cells across learning in M1 and CA3. Each point, peak response probability of a single cell from an individual session. Distributions shown separately for all recorded cells and for the subset classified as modulated, in early and late learning. Mixed-effects models accounting for repeated measurements across mice and sessions, with Benjamini–Hochberg correction across all tests: in all cells, peak response probability decreased from early to late learning in M1 (FDR-adjusted *p* < 0.001) and CA3 (FDR-adjusted *p* = 0.009). In modulated cells, peak response probability also decreased from early to late learning in M1 (FDR-adjusted *p* = 0.010); no other between-condition contrasts were significant. **B**, Fractions of excited and inhibited neurons in movement-preceded versus stationary-preceded rewarded trials at quarter-session resolution in M1 and CA3 late learning. Bars, mean fractions across quarter-session datapoints with overlaid datapoints. Classification of excited and inhibited cells as in Figure 3J. Linear mixed-effects models for movement-versus-stationary comparisons, with Benjamini–Hochberg correction across all contrasts. In M1, significant differences in the excited-fraction comparison in the Pre (FDR-adjusted *p* < 0.001) and Post (FDR-adjusted *p* < 0.001) epochs; other M1 contrasts not significant after correction. In CA3, no contrast was significant after correction. **C**, Fractions of excited and inhibited neurons during contingency-degradation (CD) sessions at quarter-session resolution in M1 and CA3, analyzed separately using reward-locked and threshold-locked alignment. Classification of excited and inhibited cells as in Figure 3J. Bars, mean quarter-level fractions with s.d. error bars; overlaid points, individual quarters. Under reward-lock alignment, modulation fractions were sparse in both regions; under threshold-lock alignment, modulation was stronger and concentrated primarily in the Pre and Post epochs, indicating that during CD modulated-cell responses remained organized around threshold crossings rather than around random reward delivery. **D–E**, Decoder performance during contingency degradation (CD) using task-modulated and non-modulated cell populations. Decoding accuracy quantified as ROC–AUC using ensembles of 10 cells per session. **D**, Threshold-to-threshold decoding (training and testing on threshold-crossing events): modulated-cell decoders outperformed non-modulated decoders (one-sided Wilcoxon, FDR-adjusted *p* = 0.005), outperformed shuffled modulated controls (FDR-adjusted *p* = 0.005), and were above chance (AUC > 0.5, FDR-adjusted *p* = 0.005). Non-modulated real decoders were not significantly above shuffled controls (FDR-adjusted *p* = 0.345) or above chance (FDR-adjusted *p* = 0.345). **E**, Threshold-to-reward transfer decoding (training on threshold-crossing events, testing on reward events): no contrast survived FDR correction after IQR filtering (modulated real vs modulated shuffle FDR-adjusted *p* = 0.345; modulated real vs non-modulated real FDR-adjusted *p* = 0.345; modulated real vs chance FDR-adjusted *p* = 0.345). Shuffled controls remained near chance (AUC ≈ 0.5) in both panels. **F–G**, PCA dimensionality of reward-manifold population activity (all recorded cells). **F**, Cumulative explained variance for M1 and CA3 in early and late learning; dashed vertical line at PC4 marks the dimensionality used in downstream low-dimensional manifold analyses. **G**, Eigenspectrum decay (mean explained-variance ratio per PC). Variance was concentrated in the first components, with progressively smaller contributions from higher-order components. Curve-level permutation tests on PCs 1–20: not significant for either metric in either region (M1 scree *p* = 0.899; M1 cumulative *p* = 0.921; CA3 scree *p* = 0.883; CA3 cumulative *p* = 0.850). Per-PC early-versus-late tests (Mann–Whitney, BH-corrected within each region and metric across PCs 1–20) yielded no significant components. **H**, Per-mouse movement-versus-stationary neural-state separation across learning in M1 (top) and CA3 (bottom). Distribution of Euclidean distances between movement and stationary trajectories at aligned timepoints in PCA state space, for individual mice, comparing early and late learning.

**Extended Data Figure 4.**
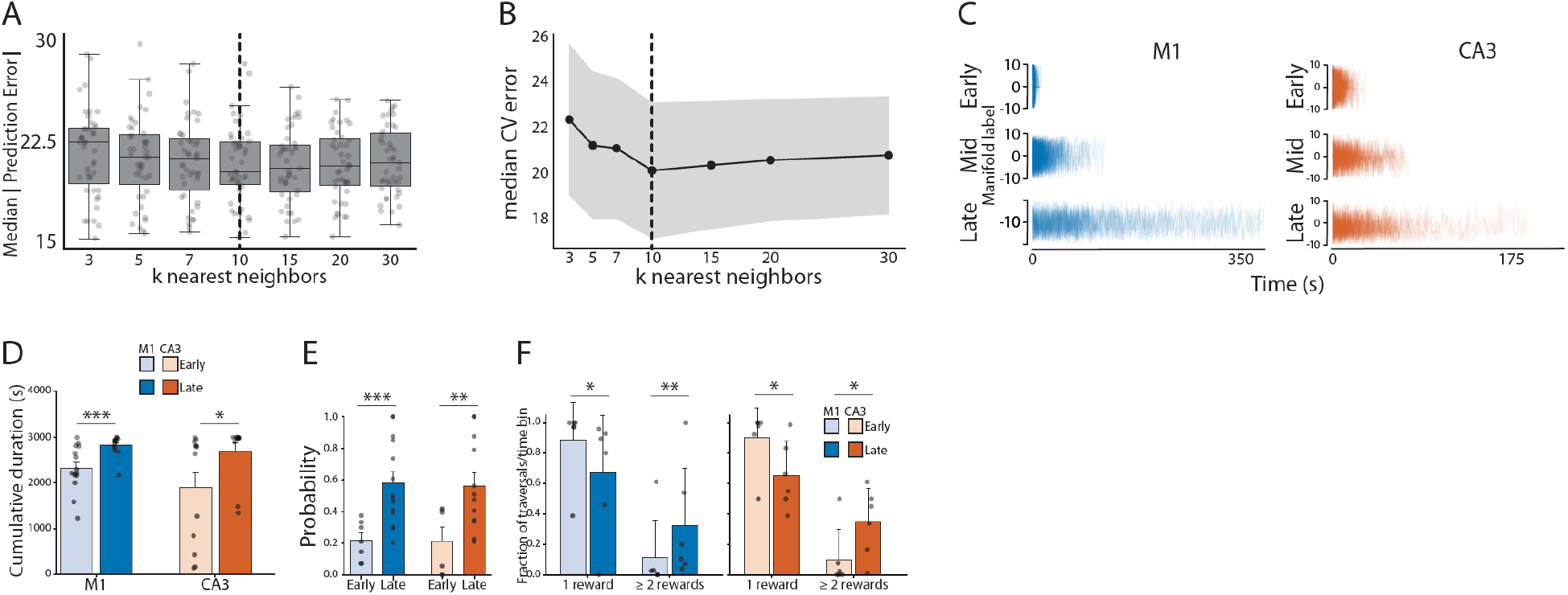
Reward-manifold traversals become increasingly structured through learning. **A–B**, Blocked cross-validation to select the number of nearest neighbors (k) for reward-manifold labeling. Blocked cross-validation performed across sessions; median absolute prediction error in reward-relative time-label bins computed for each candidate k. **A**, Session-level distributions of median prediction error for k = 3, 5, 7, 10, 15, 20, and 30 across 43 sessions. **B**, Median cross-validation error across k; shaded area, between-session variability. Lowest median error at k = 10; lower k values produced significantly higher errors than k = 10 in paired comparisons, while larger k values did not significantly improve performance, supporting use of k = 10 in the main reward-manifold analysis. **C**, Example reward-manifold traversals across learning in M1 and CA3. Representative session-wise traversals onto the reward manifold for early, mid, and late learning stages in each region. y axis, reward-relative manifold position; x axis, elapsed session time. Across learning, traversals became more prolonged and more recurrent. **D**, Total reward-manifold traversal duration per session in early and late learning for M1 and CA3. Bars, mean ± s.e.m. across sessions; overlaid points, individual sessions. Linear mixed-effects models accounting for repeated measurements across mice: significant increases in cumulative duration from early to late learning in both M1 (β = 506 s, FDR-adjusted *p* < 0.001) and CA3 (β = 762 s, FDR-adjusted *p* = 0.021). **E**, Conditional probability of reward delivery given occupancy of reward manifold traversal trajectories, P(R | reward manifold traversal). Session-level values; overlaid points, individual sessions. Linear mixed-effects models fit separately for each group, accounting for repeated measurements across mice, with Benjamini–Hochberg correction across the four tests: P(R | reward manifold traversal) increased significantly from early to late learning in both M1 (early mean = 0.216, late mean = 0.580, β = 0.364, FDR-adjusted *p* < 0.001) and CA3 (early mean = 0.213, late mean = 0.561, β = 0.348, FDR-adjusted *p* = 0.008). **F**, Fraction of rewarded traversals containing one versus multiple rewards in early and late learning for M1 and CA3. Bars, mean ± s.e.m. across mice. Paired *t*-tests: significant decrease in the fraction of single-reward traversals and corresponding increase in the fraction of multi-reward traversals from early to late learning in both M1 (*p* = 0.014) and CA3 (*p* = 0.023).

**Extended data Figure 5.**
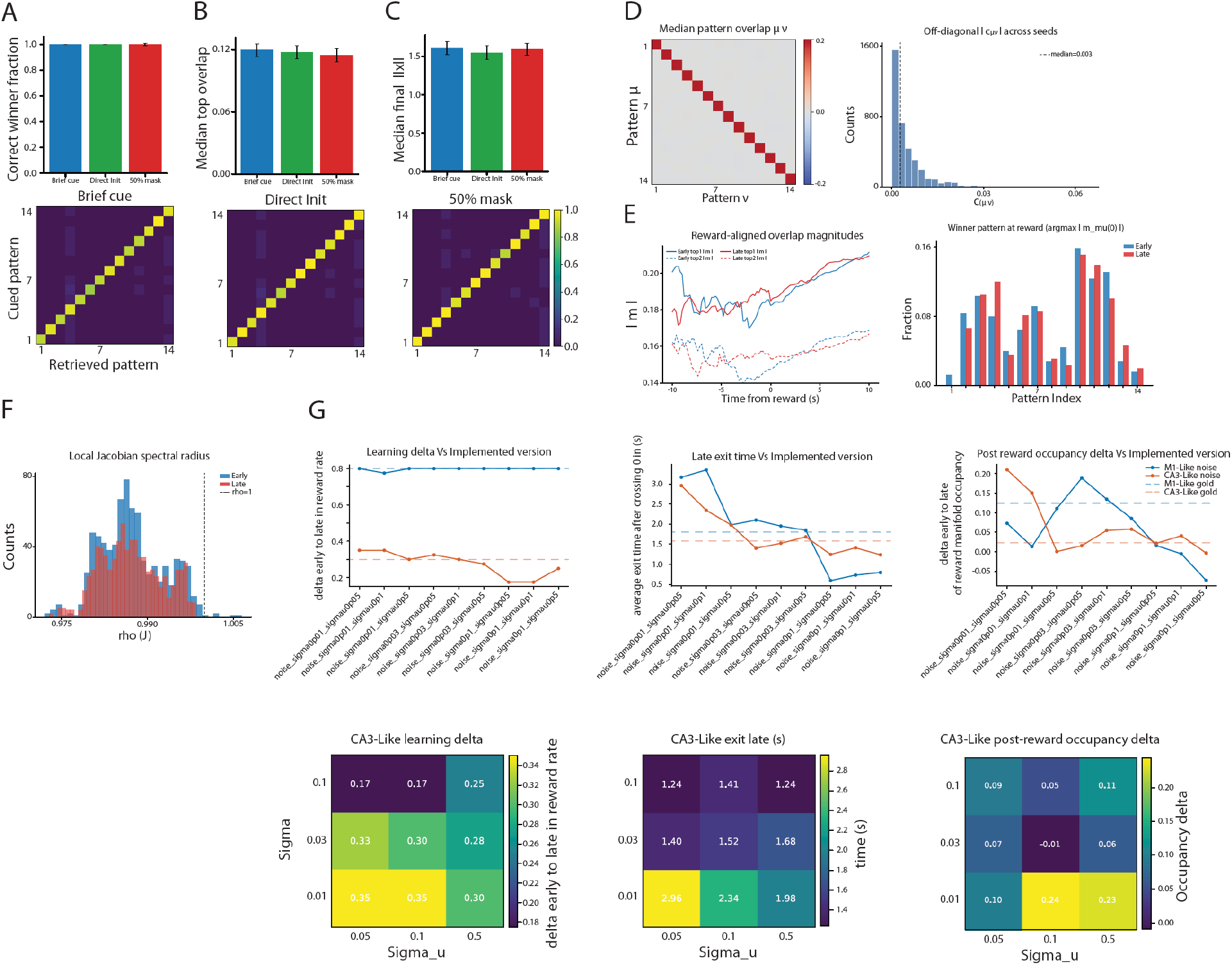
The CA3-like model exhibits robust autoassociative dynamics. **A–C**, Stored-pattern retrieval testing to verify that the CA3-like network retained an autoassociative memory structure. Fourteen stored patterns; retrieval evaluated under three perturbation regimes: brief cue (A), direct initialization from the stored pattern (B), and 50% masked cue (C). Top bar plots, retrieval accuracy, median top overlap, and final activity magnitude across seeds; lower heatmaps, mapping from cued pattern to retrieved pattern. Retrieval was strongly diagonal across all perturbation conditions, each cue returning to its corresponding stored pattern rather than converging on a single shared attractor. Median correct-winner fraction = 1.0 in all three conditions; mean retrieval accuracy = 0.925 (brief cue), 0.921 (direct initialization), 0.929 (50% mask). Median top overlap = 0.120, 0.117, 0.115; final activity magnitude = 1.61, 1.55, 1.59. **D**, Orthogonality of the stored pattern set across the 14 CA3 patterns. Left, median pattern-overlap matrix (μ, ν); strong diagonal self-overlap and near-zero off-diagonal overlap. Right, distribution of absolute off-diagonal overlap values |c_{μν}| pooled across seeds; dashed line, median = 0.003 (IQR = 0.006). Nearly orthogonal stored patterns and a low-crosstalk associative basis. **E**, Reward-aligned pattern-overlap dynamics during task performance. Left, time course of the two largest stored-pattern overlap magnitudes around reward, separately for early and late sessions; the leading overlap remained only modestly larger than the second-largest, indicating that the CA3 state remained embedded in the stored-pattern basis rather than dominated by a single fixed pattern. Median top2/top1 crosstalk ratio across seeds: 0.816 early, 0.824 late; winner margin: 0.034 early, 0.034 late. Right, distribution of winner patterns at reward; reward-period activity was distributed across multiple stored patterns rather than restricted to one. **F**, Local stability assessed by sampling the local Jacobian spectral radius along early and late CA3 trajectories. Median spectral radius similar across learning (0.988 early, 0.988 late); nearly all sampled states below the instability boundary (ρ < 1: 98.9% early, 99.9% late), indicating locally contracting dynamics around task trajectories. **G**, Robustness of the main CA3-like findings across a grid of training noise (σ) and input noise (σ_u) settings. Top, line plots of three metrics across perturbed implementations: learning delta (left), late exit time (middle), and post-reward occupancy delta (right). Each x-axis tick is a specific (σ, σ_u) combination. M1-like, blue; CA3-like, orange; dashed horizontal lines, reference (“gold”) values from the main implementation. Bottom, heatmaps of CA3-like learning delta (left), CA3-like late exit time (middle), and post-reward occupancy delta (right) across the full σ × σ_u grid (σ ∈ {0.01, 0.03, 0.1}; σ_u ∈ {0.05, 0.1, 0.5}). Core qualitative behavior preserved across perturbations: M1-like models continued to show larger learning deltas than CA3-like models, and CA3-like models retained learning across all tested noise conditions. CA3 learning delta ranged from 0.17 to 0.35 across the grid; CA3 late exit times ranged from 1.24 s to 2.96 s (reference run: 2.43 s); post-reward occupancy changes remained broadly comparable to reference values.

**Extended Data Figure 6.**
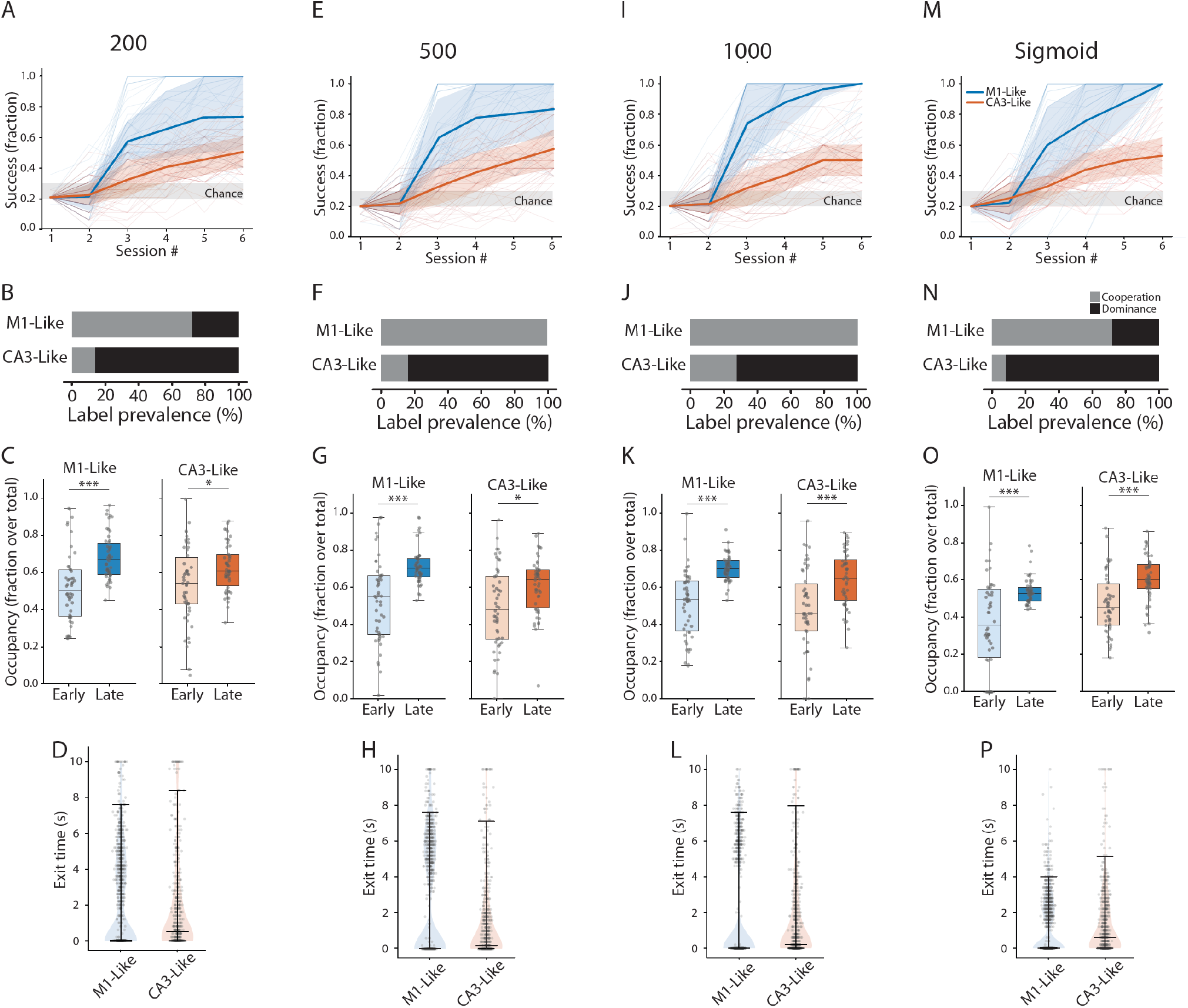
Architecture-dependent learning dynamics are robust across model implementations. A–D, E–H, and I–L show 50-seed simulations of the M1-like and CA3-like models with network sizes of 200, 500, and 1,000 units, respectively. M–P show a 50-seed sigmoid-activation control run using the same analysis format. **A, E, I, M**, Learning curves across six sessions. Thin lines, individual seeds; thick lines, median across seeds; shading, interquartile range. Gray band, chance performance (0.2). Across all network sizes and in the sigmoid-control run, both models learned above baseline, with stronger late performance in M1-like models. **B, F, J, N**, Late control-regime prevalence, summarized from reward-time positive readout-unit recruitment. Gray, cooperative recruitment of both positive readout units; black, dominant single-unit recruitment. Across the network-size sweep, CA3-like models remained biased toward single-unit dominance, whereas M1-like models showed greater cooperative recruitment, particularly at larger network sizes. The sigmoid-control run preserved the same qualitative dissociation. **C, G, K, O**, Reward-manifold occupancy from early to late learning. Occupancy, fraction of valid post-reward reward-manifold labels per seed. Paired sign-flip permutation tests, Bonferroni-corrected, significant early-to-late increases for both models across all variants: N = 200, M1-like *p* = 6.0 × 10^−5^, CA3-like *p* = 0.040; N = 500, M1-like *p* < 0.001, CA3-like *p* = 0.036; N = 1,000, M1-like *p* < 0.001, CA3-like *p* < 0.001; sigmoid control, M1-like *p* < 0.001, CA3-like *p* < 0.001. **D, H, L, P**, Late rewarded-returning exit-time distributions. Exit time, first post-reward time at which the reward-manifold trajectory exited the rewarded state. Violin plots, event-level distributions; overlaid points and summary bars. Seed-level paired comparison between M1-like and CA3-like exit times: significant for N = 200 (*p* = 0.042) and N = 500 (*p* = 0.004); not significant for N = 1,000 (*p* = 0.072) or sigmoid control (*p* = 0.500).

## Materials and Methods

### Animals

All animal experiments were approved by the Cantonal Veterinary Office Basel-Stadt, and all procedures were performed in compliance with its guidelines and the Swiss Veterinary Law (permit number 3037). This study made use of C57BL/6JRj wild type (WT) mice. Animals were housed in temperature- and humidity-controlled cages at 21–23 °C and 50–60% humidity, with 12 h – 12 h reversed light-dark cycle, and with food and water provided ad libitum expect than during behavioral training, when mice were water restricted and maintained at approximately 85% of their pre-restriction body weight, with supplemental water provided as needed after sessions. All mice were group-housed with littermates and then single-housed either after surgical procedures, or 1-3 days before the onset of behavioral experiments. For experimental procedures, mice of both sexes were randomly and blindly allocated to experimental groups by the responsible researcher.

### Surgical procedures and chronic optical access

Adult mice (> P60) were pre-treated with buprenorphine (Bupaq P, 0.1 mg/kg, s.c., Streuli) and atropine sulfate (0.05 mg/kg, s.c., Amino AG) 30 min before surgery. Anesthesia was induced with 5% isoflurane in oxygen and maintained at 1.7–2% (Attane, 0.6 L/min airflow; Piramal). A local anesthetic mix of lidocaine (< 7 mg/kg, Streuli) and bupivacaine (1–5 mg/kg, Sintetica) was administered subcutaneously over the surgical site. Mice were placed in a stereotaxic frame (1900, Kopf), and body temperature was continuously monitored and maintained using a heating pad. The scalp was sterilized with betadine (Mundipharma), and a midline incision was made. The periosteum was removed, and the skull was cleaned with betadine and sterile saline (B. Braun Surgical). The skull was scored with a needle to improve adhesion, followed by application of Histoacryl (B. Braun). Craniotomies (0.4 mm diameter drill bit, Jota AG) were drilled above the target regions. Mice received injections of pAAV.Syn.GCaMP6f.WPRE.SV40 (Addgene viral prep # 100837-AAV1; titer: 7.4 × 10^12^ or 1.3 × 10^13^ vg/mL. pAAV.Syn.GCaMP6f.WPRE.SV40 was a gift from Douglas Kim & GENIE Project^74^) into the dorsal hippocampus (AP: –1.75 mm; ML: ±1.95 mm; DV: –1.85 mm; 350 nL per hemisphere) using a Nanoject III (Drummond). Glass windows or microendoscopes (Prism-attached GRIN lenses) were lowered adjacent to the area under investigation (as in^47–48^). For M1 experiments, optical access was established through a cranial window centered on the following coordinates: Anterior-Posterior: +1.25mm from Bregma; Medio-Lateral: 1.25mm from the medial sinus. For CA3 experiments, optical access was obtained using a GRIN lens/prism-based preparation centered on the following coordinates: Anterior-Posterior: −1.58mm from Bregma; Medio-Lateral: 2.00mm from the medial sinus; Dorso-Ventral: 2.3mm from surface dura. Custom microendoscopes specifications were as follows: GRINtech microendoscope: singlet tubular GRIN lens, diameter 1.0 mm, wavelength 520nm, non-coated; prism: 1.00 × 1.00 × 1.00 mm, aluminum coating on hypotenuse. The total length of the microendoscope was approximately 4.67 mm, working distance 0.15 mm in water from prism exit surface (NA 0.4), 0.08 mm in air on the image side (NA 0.5). GRINtech GmbH, Jena. Inscopix microendoscope: cat. Nr. 1050-004601, approximate microendoscope length: 4.3 mm. The glass window or microendoscope were fixed with Venus and superglue (Loctite 415), and the surrounding skull was covered with dental cement (Paladur, Kulzer) mixed with graphite (Carl Roth, 7614). A custom metal head bar was affixed. The microendoscope was temporarily protected with Kwik-Cast (World Precision Instruments). Following recovery, mice were habituated to head fixation and to the behavioral apparatus before the start of BCI training.

### Two-photon calcium imaging

Two-photon calcium imaging was performed with a resonant-scanning B-Scope (Thorlabs) controlled by ThorImage (v4.0). Imaging was acquired at 30 Hz over a 400 µm × 400 µm field of view sampled at 512 × 512 pixels. The same field of view was followed longitudinally across training days whenever possible. Offline extraction of regions of interest (ROIs) and fluorescence time courses was performed using Suite2p^75^.

### Habituation and selection of BCI neurons on Day 0

Mice were habituated to the setup over 5 days, progressing from short head-fixation exposures to sustained sessions. After habituation, each mouse underwent a Day-0 imaging session used to identify a stable imaging plane for longitudinal tracking. Day-0 recordings were segmented with Suite2p to obtain ROI masks and fluorescence traces. Four BCI ROIs were then selected from the Day-0 segmentation: two “positive” neurons whose activity drove the decoder toward reward and two “negative” neurons whose activity opposed reward. Selection was based on spatial separation (ROIs distance between 50–150 µm), and on baseline activity (at least 10 calcium events per hour), without additional functional screening criteria. A Day-0 ROI mask was saved and used as a reference for subsequent alignment across days.

### Longitudinal field-of-view alignment and activity rate daily calibration for threshold setting

At the beginning of each training day, a short alignment recording of approximately 1–5 min was acquired to recover the Day-0 field of view. Alignment was guided by visual overlay of Day-0 ROI contours developed in Open-CaBCI which provides the live imaging image and enables manual x–y translation to achieve alignment. Before each BCI session, mice underwent a 15-min calibration recording of spontaneous activity. This calibration period was used to set a session-specific reward threshold, chosen so that spontaneous threshold crossings remained rare and chance performance was low, typically targeting approximately one random reward every 2 min, corresponding to an estimated chance reward probability of about 20–25%. The calibration procedure therefore normalized for day-to-day variability in fluorescence amplitude and spontaneous ensemble fluctuations while preserving the same causal ensemble definition across days.

### Closed-loop BCI task

Mice were trained in a trial-structured, closed-loop operant task in which reward delivery depended on moment-to-moment calcium activity of the four selected BCI neurons. The online decoder grouped the four neurons into two ensembles, with ensemble activities defined as

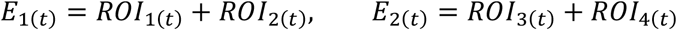

and the one-dimensional control signal (“cursor” or ensemble state) defined as

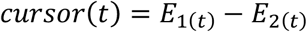

This push–pull organization discouraged nonspecific global activation and instead required differential control over the selected BCI neurons. The cursor was transformed online into an auditory feedback signal, with cursor values mapped onto discrete tone states separated by quarter-octave increments. The real-time user interface displayed reward count, auditory-feedback frequency, online ensemble state relative to threshold, fluorescence activity of the selected neurons, contour overlays, motion-correction and x–y drift information, and controls for threshold setting and contingency-degradation manipulation.

Each training day consisted of an alignment period, a 15-min calibration epoch, and a 50-min BCI session. In the BCI session, trials were structured around a 30-s response window. A trial was scored as a hit when the cursor crossed the session-specific threshold, triggering sucrose-water reward delivery. Following reward, a 3-s post-reward lockout was imposed during which additional threshold crossings were ignored and the tone was turned off. To prevent repeated rewards arising from a single prolonged excursion, a new trial was initiated only after the cursor returned sufficiently close to baseline (50% of threshold). If threshold was not reached within 30 s, the trial was classified as a miss and followed by a longer timeout implemented as a white-noise period. This task structure was maintained across longitudinal training, with an initial Day-0 selection session followed by eight training sessions in both M1 and CA3 cohorts. Mice were excluded from further analysis if their performance did not differ significantly from chance in at least two of the final three training sessions.

### Contingency degradation

In a subset of experiments, contingency degradation sessions were introduced after initial learning. During these sessions, reward delivery was decoupled from online neural control and instead delivered at an approximately matched random rate. Ensemble state threshold crossing triggered trial reset, in a similar way as regular training. This manipulation preserved overall reward statistics while disrupting the causal contingency between ensemble-state threshold crossing and reward. Recovery sessions were then acquired after contingency degradation.

### Online fluorescence processing

For each imaging frame, fluorescence within each ROI was computed by summing pixel intensities inside the ROI and normalizing by ROI size. Fluorescence was converted online to ΔF/F using a baseline F_0 that was updated periodically, typically every 120 s, to compensate for slow drifts. To reduce noise while maintaining low latency, ROI signals were smoothed online by averaging over 6 consecutive frames, corresponding to 0.2 s at 30 Hz. The smoothed ROI values were then entered into the ensemble-state computation described above.

### Behavioral monitoring and movement-based trial classification

Behavior was monitored with a live video feed and wheel rotation was measured using a rotary encoder. For analyses that separated trials into movement-preceded and stationary-preceded categories, wheel-velocity segments aligned to reward were examined and three velocity-derived criteria were combined using a logical AND rule. Specifically, trials were labeled “move” only when sustained-movement, peak-velocity and mean-velocity criteria were all satisfied; all other rewarded trials were labeled “stay” or stationary-preceded. Example locomotor traces aligned to trial onset were used as a quality check to confirm that the two behavioral classes were cleanly separated in the expected pre-trial time window. This strict criterion was intended to favor specificity over sensitivity, thereby producing conservative movement labels for downstream neural-state geometry analyses.

### Offline calcium preprocessing

Offline fluorescence traces were extracted from Suite2p output. For manifold and population-state analyses, detrended calcium traces were converted into sparse binary event rasters using a two-step procedure. First, each trace was smoothed by averaging within non-overlapping blocks of 7 frames, corresponding to 233 ms at 30 Hz, and the block mean was reassigned to all 7 frames. Second, event onset was detected using a threshold-plus-rising-edge rule: the signal had to exceed threshold and its first difference had to be positive. Once detected, the event vector was set to 1 for the following 7 frames, thereby imposing a minimum event width and merging closely spaced crossings. Unless otherwise noted, the event threshold was 0.05 ΔF/F; for a small number of sessions, a lower threshold of 0.025 was used according to session-specific criteria. For state-space analyses, event rasters were further down sampled by grouping 6 consecutive imaging frames into one bin, yielding an effective sampling rate of 5 Hz (0.2 s per bin). Reward times recorded in imaging frames were converted to this same binned time base for alignment.

### Reconstruction of trials and reward times offline

To standardize trial-based analyses across figures, task structure was reconstructed offline from the stored cursor trace, threshold and timing data stored. Reward times were identified as threshold crossings subject to the same post-reward lockout used online. Miss trials were defined by failure to reach threshold within the allowed trial duration and were followed by an additional lockout corresponding to the white-noise timeout. This reconstruction produced reward times, trial boundaries, and timeout periods for each session, allowing all downstream performance, geometry and peri-reward analyses to be computed from a common inferred trial structure. Ensemble-trajectory directness was computed as the net displacement from trial start to trial end divided by the cumulative absolute displacement across the trajectory.

### Performance measures

Behavioral performance was quantified primarily as hit rate, defined within each 5-min bin as the number of rewarded trials divided by the number of trial starts. Additional measures included within-session reward rate, rewarded-trial duration, threshold-crossing rate, the fraction of successful movement-preceded versus stationary-preceded trials, and trajectory directedness in tone or ensemble-state space.

Directedness was computed for each trial segment as signed path efficiency:

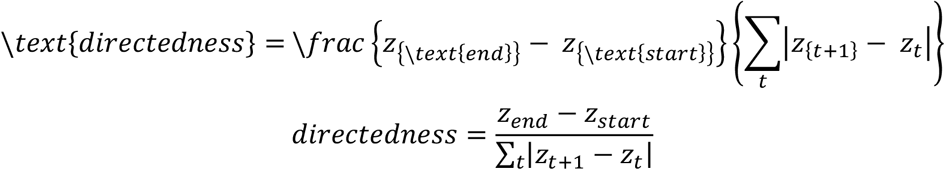

where z_t denotes the tone or ensemble-state value over the trial interval, typically from trial start to trial end as defined in the trials table. Values near +1 indicate a highly directed increase, values near −1 indicate a highly directed decrease, and values near 0 indicate tortuous or non-directed trajectories with little net displacement relative to total path length. Trial-level directedness values were averaged at the session level or within 5-min bins, depending on the figure. For early-versus-late comparisons, early learning was generally defined as the first two valid training sessions and late learning as the last two valid training sessions; manipulation sessions, including CD and REC, were analyzed as separate phases.

### Reward-triggered analyses of BCI neurons and population activity

For direct-neuron and ensemble analyses, ROI traces and ensemble signals (E_1, E_2, and cursor) were aligned to reward delivery and averaged across rewarded trials within session. These reward-triggered signals were summarized using peak-response measures and changes relative to pre-reward baseline. At the network level, reward-triggered burst probability time courses were computed by extracting the binarized event trace of each neuron in a ±10 s window around each reward and averaging across rewards. These peri-reward profiles formed the basis for identifying task-modulated neurons and for quantifying excitation–inhibition balance across learning.

### Identification of reward-modulated neurons

To identify reward-modulated cells, each neuron’s reward-triggered time course was divided into four predefined temporal epochs: Long-Pre (−10 to −3 s), Pre (−3 to 0 s), Post (0 to 3 s), and Long-Post(3 to 10 s)^22^. Within each epoch, a linear trend was fit and Pearson correlation with time was computed. A neuron was classified as modulated if, in at least one epoch, the absolute slope exceeded 10^−3^ and the corresponding significance criterion satisfied p<0.001. This was implemented as an OR rule across epochs. Within each epoch, a modulated neuron was classified as excitatory if the trend was significantly positive and inhibitory if the trend was significantly negative. Fractions of excited and inhibited neurons were then summarized per session, per animal, or per quarter-session depending on the analysis. The fraction of modulated neurons,

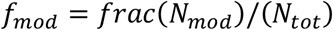

as used as a network-sparsification measure. This same slope/correlation/significance framework was extended to reward-lock, threshold-lock, movement-preceded and stationary-preceded analyses.

### Longitudinal Rank-Shift Analysis of Tracked Cell Activity Across Learning

As an auxiliary longitudinal analysis, we quantified changes in activity ranking among cells that could be tracked across learning. For each animal, cells were identified and analyzed longitudinally across training days. For each tracked cell, we extracted a day-wise activity-strength proxy from the average trace by taking the within-day maximum, yielding a day-by-cell matrix of activity scores.

Training sessions were ordered chronologically and split into early and late learning windows, corresponding to the first sessions versus the last sessions. For each cell, we computed mean activity strength in the early and late windows and converted these values into within-window rank positions across the tracked population.

We then computed a rank-shift metric, defined as the late rank minus the early rank, with the sign interpreted according to the ranking convention. Rank shifts were summarized for modulated cells in M1 and CA3. This analysis was used as a secondary measure of longitudinal reorganization and was not part of the core reward-manifold pipeline unless explicitly shown in the corresponding figure panels.

### Decoding and cross-day stability analyses

Task-modulated neurons were also used for decoder-based and longitudinal identity analyses. Decoder performance was quantified using ROC–AUC, typically comparing the top modulated cells with equally sized control populations of randomly selected non-modulated cells and shuffled-label baselines. During contingency degradation, both threshold-to-threshold and threshold-to-reward decoding configurations were examined. Cross-day stability of modulated-cell identity was quantified using Jaccard similarity. Rank-shift analyses quantified learning-related reordering of modulated-cell prominence across training. These analyses were performed at mouse level after cell identity had been tracked across days through the longitudinal field-of-view alignment procedure. Decoder performance was quantified with L2-regularized logistic regression (liblinear, class-balanced) using blocked temporal cross-validation; ROC–AUC was averaged across held-out folds. We compared top modulated-cell decoders to equal-size random non-modulated controls and shuffled-label nulls (N shuffles). During contingency degradation we evaluated threshold-label and reward-label decoding configurations using the same pipeline. Cross-day stability of modulated-cell identity was quantified by Jaccard similarity of session-wise modulated-cell sets, and learning-related reordering was quantified by rank shifts of modulation strength across early vs late sessions after longitudinal cell registration.

### PCA-Based Quantification of Dimensionality

To quantify the dimensional structure of the population activity underlying the population analysis, we applied principal component analysis (PCA) to the same session-wise activity matrices used for manifold construction. We built, for each session, a neuron-by-time activity matrix from reward-aligned population traces (same alignment window, binning, and preprocessing used for the manifold analyses). Rows corresponded to neurons and columns to time bins (or trial-averaged time bins, depending on panel). This matrix was mean-centered across time per neuron, and PCA was then applied to its transposed form (samples = time bins/trial-time points, features = neurons). For each session, we extracted the PCA eigenspectrum and cumulative variance explained, allowing us to estimate how rapidly population variance was captured by successive components.

To control for differences in population size, we repeated the same PCA procedure using a fixed-size subset of the top 20 valid neurons per session; when fewer than 20 valid neurons were available, all available valid neurons were retained. This provided a neuron-count–matched dimensionality estimate for comparison with the full-population analysis.

The variance explained by each principal component was computed from the PCA eigenvalues and normalized by the total variance, while cumulative variance was obtained as the running sum across ordered components. A shared low-dimensional embedding was fit jointly by concatenating the time samples from the included conditions and computing PCA, using the first four principal components, which were enough to capture >50% of the population variance (all subsequent components individually contributed ≤5% additional explained variance).

Sessions were grouped by brain region and learning phase, yielding four groups: M1 early, M1 late, CA3 early, and CA3 late. For each group, we summarized the explained variance at each component as the mean ± s.e.m. across sessions. Scree curves were used to compare the decay of variance across principal components, whereas cumulative variance curves quantified the fraction of total population structure retained by low-dimensional embeddings, including the variance captured by the first four components.

To assess whether these estimates were driven by differences in cell number or cell selection, we repeated the same analysis using a top-20-neuron fixed-size control. This control tested whether group differences in variance retention were preserved under a constrained population size, while allowing sessions with fewer than 20 valid neurons to contribute all available valid neurons. Using this top-20 fixed-size control, the same qualitative pattern was preserved: CA3 retained variance more quickly than M1 (steeper cumulative curves and faster scree decay), and no clear early-versus-late shift was detected within region. Thus, the dimensionality differences were robust to neuron-count matching and were not explained by unequal population size.

Together, these analyses quantified how much of the reward-manifold population structure was retained in low-dimensional space and allowed direct comparison of dimensionality across region, learning phase, and analysis variant.

### Population-state geometry and trial-type representation analysis

To quantify how learning reshaped trial-type representations, high-dimensional population response vectors were assembled from trial-aligned activity, usually restricted to the subset of reward-modulated neurons to emphasize task-relevant population dimensions. Two event classes were analyzed: reward delivery and wheel rest-to-movement transitions. For each event at time t_0, a peri-event window of ±10 s was extracted, and a narrower analysis window, typically −5 to 5 s, was used for state-space geometry. Trials were split into movement and stationary using the behavioral criteria described earlier, and trial-averaged responses were computed separately for each condition. To stabilize stage-level estimates, trial-averaged responses were further averaged across two early and two late sessions. Wheel-onset events within ±5 s of reward were excluded from wheel-triggered analyses to reduce contamination by reward-related activity.

For dimensionality reduction, a shared low-dimensional embedding was fit jointly across all trajectories by concatenating the time samples from the included conditions and computing PCA, using the first four principal components, which were enough to capture >50% of the population variance (all subsequent components individually contributed ≤5% additional explained variance). Missing values were forward-filled within trajectory segments when needed. Neural-state distance was defined as Euclidean distance between condition-matched points in the common embedding at each aligned time point. Distances were then summarized across the predefined analysis window by mean or area under the distance curve to avoid multiple-comparison inflation across time. These metrics were used to quantify separation between movement and stationary trajectories within early and late learning, as well as deformation between early and late trajectories within a given class.

### Learning-Related Reorganization of Movement and Stationary Population Geometry

To quantify how learning reshaped population-level geometry associated with movement and stationary behavioral states, we analyzed longitudinal changes in PCA trajectory structure at the level of individual mice. For each mouse, we used merged PCA containing movement- and stay-aligned population trajectories. Analyses were performed separately by brain region and learning phase, allowing us to compare how M1 and CA3 reorganized movement and stationary state representations across training.

In the baseline analysis, we first quantified the separation between movement and stationary trajectories during early and late learning. For each mouse, trial-wise movement-versus-stationary distances were computed within each epoch, and learning-related change was defined as the difference between late and early mean separation. Mice were classified as showing either divergence or convergence depending on whether movement-stationary separation increased or decreased across learning. These patterns were visualized as per-mouse early–late distributions, grouped by trend and region.

We next asked whether learning preferentially shifted movement or stationary trajectories. To do this, we compared early-to-late distances within each modality: movement trajectories were compared between early and late learning, and stationary trajectories were compared across the same epochs. For each mouse, we then determined whether the movement-related shift exceeded the stationary shift, or vice versa. Region-level summaries were represented as the proportion of mice showing stronger motor versus stationary manifold reorganization.

To further assess the stability and evolution of trajectory geometry within sessions, we performed a half-session modality analysis. Each session was divided into two equal trial halves, separately for movement and stationary trials. Population trajectories from each half were projected into a common PCA space, and Euclidean distances between half-session trajectories were computed across early and late learning.

Phase effects were tested using linear mixed-effects models with learning phase as a fixed effect and mouse as a random intercept. P-values were corrected across panel-wise tests using Benjamini–Hochberg false-discovery-rate correction.

Finally, we summarized the directionality of learning-related geometric change by taking the sign of the late-minus-early change in movement-stationary separation for each mouse. This produced a binary classification of mice showing increased versus decreased movement–stationary separation with learning. The distribution of these signs was compared between regions using Fisher’s exact test, providing a region-level test of whether M1 and CA3 differed in the direction of representational change.

To test whether these geometric effects depended on the number or identity of neurons included in the analysis, we repeated the full pipeline using top-k restricted populations. Specifically, we generated matched analyses using the top 10, top 20, and top 50 modulated cells per mouse. Cells were ranked according to modulation strength, defined as the session-averaged absolute difference between movement and stationary mean responses within the analysis window.

For each mouse, the k most strongly modulated valid cells were retained. When fewer than k valid cells were available, all available cells were used. Top-k analyses were implemented using both precomputed top-k manifold files and direct re-computation where required, with consistent downstream distance metrics, visualizations, and statistical tests applied across variants.

Together, these analyses quantified how movement and stationary population trajectories changed across learning, whether these changes reflected divergence or convergence between behavioral states, and whether the observed region-specific geometry was robust to controlled neuron-count constraints.

### Nonparametric reward-manifold labeling and traversal analysis

To quantify ongoing access to reward-aligned subspaces without fitting an explicit latent-variable model, a nonparametric recurrence-based manifold-labeling procedure was developed. At each 5-Hz time bin, population activity exclusively from modulated neurons was represented as a binary vector x_t. Reward times were first used only to initialize labels: for each reward, bins spanning 10 s before to 10 s after reward were assigned integer offsets relative to reward, with negative values for pre-reward bins, 0 at reward, and positive values for post-reward bins; all other bins remained unlabeled (NaN). Euclidean distances were then computed between population vectors, and for each query time point the k = 10 nearest neighbors were identified after excluding candidate neighbors within ±10 s of the query time to eliminate trivial local autocorrelation. The median reward-relative label of these neighbors, ignoring NaNs, was then assigned to the query bin and converted from bins to seconds. If all nearest-neighbor labels were NaN, the query bin remained undefined. The resulting one-dimensional reward-manifold coordinate was smoothed with a 3-bin NaN-aware moving average, corresponding to approximately 0.6 s.

Contiguous stretches of non-NaN manifold coordinate were defined as reward-manifold traversals. Traversal duration was the segment length multiplied by 0.2 s. A traversal was labeled reward-associated if at least one reward occurred within its temporal bounds; otherwise, it was classified as non-rewarded. Session-level measures included the number of traversals, total time spent in labeled states, the number of zero crossings across the reward-centered boundary, the fraction of rewarded traversals, the conditional reward probability inside versus outside classified states, the fraction of rewarded traversals containing one versus multiple rewards, and the post-reward occupancy of positive manifold states.

Trace-based crossings were defined as sign changes of the continuous reward-manifold coordinate across 0, whereas neighbor-label-based crossings were defined as time points whose nearest-neighbor label set included the reward-centered label. Rewarded crossings analyses were event-aligned rather than full-trial endpoint relabeling analyses. Specifically, after computing the continuous manifold-state value at every time bin, zero-crossing events were detected, and only crossings occurring near the reward were retained. Windows were then extracted around these crossing times and averaged by group and learning phase. Thus, the labels are reward-based, whereas the PSTHs are crossing-based: they quantify local neural-state dynamics around reward-coupled crossings, not monotonic progression from trial start to trial end. Under this procedure, a non-monotonic PSTH indicates that trajectories tend to approach the reward-centered boundary and then return, rather than continuously passing through it. This framework allowed learning-dependent changes in recurrence, occupancy, reward association, and traversal directionality to be quantified at the session, animal, and traversal levels.

To make `k` selection explicit, we performed blocked time-series cross-validation on the same reward-manifold traversal sessions used in the main analysis. For each candidate `k`, we predicted held-out frame time-to-reward labels from k-nearest neural population states, while excluding neighbors within a temporal lockout window to reduce autocorrelation leakage. We evaluated median absolute prediction error per session and compared candidates at the session level. This analysis shows where `k=10` lies relative to the cross-validated optimum and whether nearby alternatives produce meaningfully lower error. Best k by median blocked-CV absolute error is k=10 (43 sessions).

Median CV error (time-label bins): k=10: 20.10 (lowest); k=3: 22.33; k=5: 21.20; k=7: 21.07; k=15: 20.33; k=20: 20.55; k=30: 20.77. Paired session-wise Wilcoxon vs k=10: k=3, k=5, k=7 are significantly worse (p=0.0010, 0.019, 0.037); k=15,20,30 not significantly different from k=10.

### Traversal-Level Quantification of Reward-Manifold State-Space Kinematics

To characterize the dynamical structure of reward-manifold trajectories, we performed state-space analyses at the level of individual traversals. Traversals were grouped according to brain region and learning phase, yielding four conditions: M1 early, M1 late, CA3 early, and CA3 late. This allowed us to compare how population trajectories explored and moved through the reward-related manifold across learning.

For the joint position–velocity analysis, each traversal trajectory was represented as a one-dimensional position trace, x(t), along the reward-manifold coordinate. In the primary formulation, velocity was estimated using finite differences between consecutive samples and paired with the corresponding midpoint position. Position–velocity samples were then pooled within each condition to visualize the occupancy of trajectories in (x, v) space. These joint histograms provided a phase-space view of traversal dynamics, capturing both where trajectories were located on the manifold and how rapidly they moved through those positions. Central-difference variants were also performed as sensitivity controls to verify that the observed structure was not an artifact of the forward-difference midpoint transform.

To quantify the extent of state-space exploration, rewarded-returning traversals were projected onto a common two-dimensional grid defined over the pooled position and velocity ranges. The grid consisted of 60 × 60 bins spanning percentile-bounded (x, v) limits, reducing the influence of extreme values while preserving the shared dynamical range across conditions. For each traversal, we computed the fraction of explored space as the number of unique grid bins visited divided by the total number of bins.

All available traversals were retained in the primary analysis. Region differences within each learning phase were assessed using mouse-cluster permutation tests, in which group labels were shuffled at the mouse level while preserving the nested traversal structure within each animal. Where multiple phase-wise comparisons were performed, p-values were corrected using Benjamini–Hochberg false-discovery-rate correction.

We next asked whether trajectories slowed near the center of the manifold. For each condition, near-center and far-from-center spatial bands were defined from pooled quantiles of absolute manifold position within that condition. For each traversal, we computed the median absolute velocity within the near and far bands and summarized near-center slowing as the ratio between median near velocity and median far velocity. Ratios below one therefore indicate reduced velocity near the manifold center.

Within each condition, near-versus-far velocity differences were tested using one-sided Wilcoxon signed-rank tests, with the alternative hypothesis that velocity was lower near the center. Between-region differences in slowdown ratio were tested separately within early and late learning using two-sided Mann–Whitney U tests.

Finally, we quantified the temporal dispersion of traversal trajectories using mean-squared displacement. For each traversal, mean-squared displacement was computed as a function of temporal lag using fixed lag bins and truncated to a common maximum lag, including the 0–3 s window used for the final comparisons. Condition-level MSD curves were summarized as mean ± s.e.m. across traversals.

In addition, scalar MSD summaries were extracted at the terminal lag of the selected window and compared across predefined contrasts using Mann–Whitney U tests. To complement these scalar comparisons, we also performed curve-level permutation analyses on lag-resolved MSD differences, allowing us to test whether early and late learning differed in their full dynamical profiles rather than only at a single time lag.

Together, these analyses quantified three complementary aspects of reward-manifold dynamics: the structure of position–velocity occupancy, the extent of explored dynamical space, and the temporal spread of trajectories. This framework allowed us to compare whether M1 and CA3 traversals occupied, traversed, and dispersed through the reward manifold differently across learning.

### In silico BCI learning in Recurrent Neural Networks (RNNs)

#### Computational framework and study design

All analyses were performed in a discrete-time, fully *in silico* closed-loop brain–computer interface (BCI) framework. We compared two recurrent network classes under the same task protocol: a low-rank cortical reference model (M1) and an associative recurrent model with symmetric connectivity (CA3). Unless otherwise stated, the reference analyses used 20 independent random seeds per model. For each seed, the task schedule, trial structure, input statistics, and optimization schedule were matched across models. The principal summary measures were session-wise reward rate, low-dimensional trajectory geometry, reward-manifold occupancy, and post-reward manifold retention.

#### State variables and readout

In both models, the latent state *x*_*t*_ ∈ ℝ^*N*^ evolved in discrete time and the observable population activity was

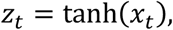

with *N* = 128 recurrent units in the reference regime. The task readout was defined from a fixed four-unit ensemble selected once per seed without replacement. Two units contributed positively and two negatively, yielding

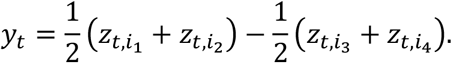

A trial was classified as rewarded at the first-time bin for which *y*_*t*_ > θ, where θ denotes the reward threshold. In the CA3 model, stored patterns were orthogonalized to the readout direction in the reference regime to reduce trivial alignment between the associative basis and the reward readout.

#### M1-like model

The M1-like model combined a fixed random recurrent backbone with a trainable rank-*r* recurrent term, where *r* = 12 in the reference regime. Its update equation was

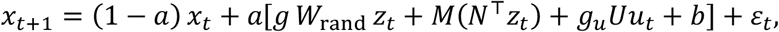

where *W*_rand_ ∈ ℝ^*N*×*N*^ is a fixed Gaussian random matrix, *M, N* ∈ ℝ^*N*×*r*^ are trainable low-rank factors, *U* ∈ ℝ^*N*×*d*^ is the trainable input matrix, *b* ∈ ℝ^*N*^ is a trainable bias, *g* scales the fixed recurrent backbone (fixed term at 0.9), *g*_*u*_ scales the input drive, and ε_*t*_ ~ 𝒩(0, *σ*^2^*I*) is additive internal noise. Each entry of the fixed random recurrent backbone was drawn independently from a Gaussian distribution with mean zero and standard deviation equal to 0.6 divided by the square root of the number of neurons. The recurrent gain was then applied multiplicatively to this fixed backbone during the dynamics. In the reference configuration, the M1 context-gated recurrence option was disabled. The retained trainable parameters were the low-rank factors *M, N*, the input matrix *U*, and the bias *b*. The intended low-rank constraint applies only to the trainable recurrent component, so the model should be described as a full-rank fixed random recurrent backbone plus a trainable rank-12 recurrent perturbation.

#### CA3-like model

The CA3-like model used a fixed symmetric associative core constructed analytically from stored patterns. Let *P* ∈ ℝ^*N*×*M*^ denote the stored-pattern matrix, with *M* = 14 in the reference regime. The recurrent operator was

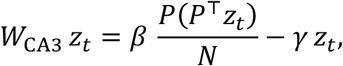

with gain parameter *β* and inhibitory stabilization *γ*. The state update was

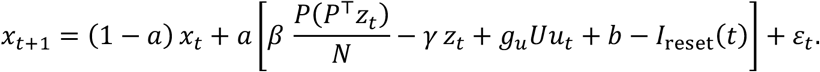

In the reference regime, *I*_reset_(*t*) was parameterized but set to zero (reset *I*_0_ = 0), so the effective CA3 dynamics were governed by the fixed associative term, the trained input/bias pathways, and additive noise. Training did not modify the recurrent associative core in the main analyses; only *U* and *b* were optimized.

#### Task protocol and inputs

Each run consisted of 6 sessions with 20 trials per session. The simulation step size was *Δt* = 0.2 s and each trial lasted 30 s, yielding 150-time bins per trial. The external input had dimension *d* = *r*. For each trial,

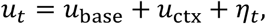

where *u*_base_ is a fixed tonic component, *u*_ctx_ is a context-dependent component, and *η*_*t*_ is Gaussian input noise. In the reference regime, the task used two orthogonal context templates corresponding to *move* and *stay*. Trial context was sampled uniformly at random. Between-trial carryover was enabled by initializing each new trial from the final latent state of the preceding trial, thereby preserving sequential dependence across trials.

#### Threshold calibration and reward definition

The reward threshold was calibrated once per seed on day 0 and then held fixed for all subsequent sessions. Calibration used 30 trials with learning frozen and estimated a threshold intended to yield approximately 20% rewarded trials at baseline. During simulated behavior, reward time was defined as the first bin at which the signed readout *y*_*t*_ crossed threshold. If no crossing occurred within the trial, the trial was unrewarded. Following rewarded trials, the model entered a post-reward wait period of up to 8 s; following unrewarded trials, it evolved for a 10 s failure timeout before the next trial.

#### Optimization objective and parameter updates

Parameters were updated with Adam using learning rate 3 × 10^−3^, *β*_1_ = 0.9, *β*_2_ = 0.999, ϵ = 10^−8^, and gradient clipping at 1.0. The leak parameter was *a* = 0.2, the internal noise standard deviation was *σ* = 0.03, and the reward-surrogate temperature was 1.0. Let

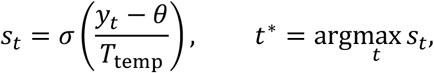

where *σ*(⋅) denotes the logistic sigmoid and *T*_temp_ the surrogate temperature. The scalar objective minimized on each trial was

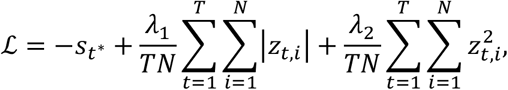

with *λ*_1_ = 10^−3^ and *λ*_2_ = 5 × 10^−4^. Thus, optimization combined a differentiable reward-surrogate term with sparse and quadratic activity penalties. Parameter updates were applied after every trial, while reward-rate summaries were reported at the session level.

#### Population trajectories and separability analysis

Population analyses were performed on the post-nonlinearity activity *z*_*t*_. For each seed, trial-averaged trajectories were computed separately for the four condition combinations

{early move, early stay, late move, late stay},

where *early* and *late* corresponded to sessions 0 and 5, respectively, unless otherwise specified. Principal component analysis (PCA) was then fit separately for each seed to the concatenation of the four condition-averaged trajectories, providing a common low-dimensional coordinate frame within seed. Move–stay separability was defined as the mean Euclidean distance between the move and stay trajectories in the retained PCA space. The reported separability change was

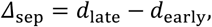

where *d*_early_ and *d*_late_ denote the time-averaged move–stay distances in early and late sessions.

Reward-manifold labeling, occupancy, and retention. Reward-manifold labels were computed from late-session activity using a *k*-nearest-neighbors transfer rule in population space with Euclidean distance and *k* = 10. First, bins within ±10 s of each reward event were assigned a base label equal to their signed time relative to reward. Labels for all remaining bins were then inferred from neighboring points in population space, excluding neighbors within the same local temporal window to avoid trivial temporal leakage. The resulting label trace was smoothed with a 3-bin NaN-aware moving average. From these labels we computed: (i) manifold occupancy, defined as the fraction of bins receiving a non-NaN reward-manifold label; (ii) the number and mean duration of contiguous occupancy epochs; (iii) crossings of the reward manifold; and (iv) post-reward manifold retention and exit time. Early and late occupancy analyses were computed separately by session.

#### Reward modulation and ensemble-drive analyses

Reward-aligned single-unit modulation was quantified from rewarded trials only. For each neuron, we computed

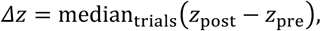

where *z*_pre_ is the mean activity in the [−5, −3] s window relative to reward and *z*_post_ is the mean activity in the [0,3] s window. A neuron was classified as modulated when |*Δz*| exceeded the analysis threshold used in the corresponding panel. Ensemble-drive analyses were evaluated at reward time from the two positive readout units. In the default imbalance criterion,

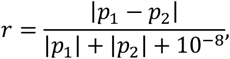

where *p*_1_ and *p*_2_ are the two positive-unit activities at the reward bin. Trials were classified as co-driving when *r* < 0.2 and as single-driving otherwise. In the alternative contribution-based criterion, positive contributions were rectified, normalized, and categorized by whether both units contributed at least 20% or a single unit exceeded 80% of the total positive contribution.

#### Hopfield-style validation of the CA3 prior

The CA3 prior (fixed symmetric autoassociative recurrent architecture used in the CA3-like model) was validated independently of task learning by testing three standard recall conditions: exact cue presentation, direct initialization at a stored pattern, and 50% masked cue completion. In these analyses, the network was initialized or briefly driven toward a stored pattern and then allowed to evolve autonomously under the fixed associative dynamics. Performance was summarized by winner accuracy, top-overlap magnitude, margin to the runner-up pattern, final state norm, and the number of distinct retrieved endpoints across patterns.

#### Sigmoid Control

To test whether the models’ performances depended on the choice of activation function, we repeated the reference simulations using a sigmoid activation function in both the M1-like and CA3-like models. All other core task and training settings were kept matched to the reference configuration, including six training sessions, 20 trials per session, day-0 calibration to a baseline reward fraction of 0.2, and the same trainable parameter sets for each architecture. The sigmoid-control analysis was run for 50 random seeds. Learning curves, late positive-unit driving labels, reward-manifold occupancy, and post-reward exit-time analyses were computed using the same pipeline as in the main model analyses.

#### Network-Size Scaling

To evaluate robustness to finite network size, we repeated the model comparison across networks with 200, 500, and 1000 units. For each network size, we ran 50 seeds for both the M1-like and CA3-like models. The M1-like model retained the same low-rank trainable component and fixed random recurrent backbone, while the CA3-like model retained the same symmetric autoassociative structure and trainable input pathways. Apart from network size, task structure, training schedule, calibration procedure, and analysis windows were kept matched across conditions. For each run, performance was summarized across six sessions as reward fraction per session. Late driving regime was quantified from the relative recruitment of the two positively weighted readout units at reward time and classified as cooperative or single-unit dominated. Reward-manifold occupancy was measured as the fraction of valid post-reward reward-manifold labels in early and late phases. Exit time was measured on rewarded-returning trials as the first post-reward time at which the reward-manifold trajectory crossed below zero. As with the main run, we restricted the analyses to rewarded-returning trajectories crossing back below zero within the +10 s post-reward analysis window.

### Statistical analysis

Unless otherwise stated, analyses were performed on session-level, animal-level, traversal-level, or 5-min-bin-level summaries as appropriate for each figure panel. Early and late learning stages were selected according to a predetermined rule, i.e. first two versus last two valid training sessions, with contingency-degradation and recovery sessions excluded from standard learning-stage comparisons. Across the paper, inferential procedures included linear mixed-effects models for repeated-measures data, Wilcoxon signed-rank tests for paired nonparametric comparisons, paired t-tests in selected matched analyses, paired sign-flip permutation tests for matched calibration-versus-session comparisons, and label-permutation tests for group or trajectory-curve comparisons. For time-resolved curves, inferential testing was performed on summary measures such as mean distance or area under the curve over a predefined time window rather than at each time point. Mixed-effects models included mouse and session as a nested variance component. Multiple comparisons were corrected using the Benjamini–Hochberg false-discovery-rate procedure across the family of tests corresponding to each panel or analysis set.

Statistical analyses were performed using Python (version 3.8). Box plots display the 25th percentile, the median, the 75th percentile, with whiskers extending to the minimum and maximum value. The box itself represents the interquartile range (IQR), encompassing the middle 50% of the data. The line within the box indicates the median. Individual data points represent either individual animals or neurons, and are shown in each figure unless otherwise stated. Statistical significance is indicated throughout the figures and legends, using the following conventions: *p < 0.05, **p < 0.01, ***p < 0.001. Graphs were generated in Python, and further edited using Adobe Illustrator (versions 2024 and 2025). While no statistical methods were used to predetermine sample sizes, all sample sizes were consistent with those reported in previous studies, and post hoc effect sizes were computed and reported.

